# Phosphorylation of pyruvate dehydrogenase marks the inhibition of *in vivo* neuronal activity

**DOI:** 10.1101/2023.03.13.532494

**Authors:** Dong Yang, Yu Wang, Tianbo Qi, Xi Zhang, Leyao Shen, Jingrui Ma, Zhengyuan Pang, Neeraj K. Lal, Daniel B. McClatchy, Kristina Wang, Yi Xie, Filip Polli, Anton Maximov, Vineet Augustine, Hollis T. Cline, John R. Yates, Li Ye

## Abstract

For decades, the expression of immediate early genes (IEGs) such as c-*fos* has been the most widely used molecular marker representing neuronal activation. However, to date, there is no equivalent surrogate available for the decrease of neuronal activity (i.e., inhibition). Here, we developed an optogenetic-based biochemical screen in which population neural activities can be controlled by light with single action potential precision, followed by unbiased phosphoproteomic profiling. We identified that the phosphorylation of pyruvate dehydrogenase (pPDH) inversely correlated with the intensity of action potential firing in primary neurons. In *in vivo* mouse models, monoclonal antibody-based pPDH immunostaining detected neuronal inhibition across the brain induced by a wide range of factors including general anesthesia, sensory experiences, and natural behaviors. Thus, as an *in vivo* marker for neuronal inhibition, pPDH can be used together with IEGs or other cell-type markers to profile and identify bi-directional neural dynamics induced by experiences or behaviors.

## Introduction

In the 1980s, it was discovered that transcription of the c-*fos* gene was robustly induced by neuronal activity^1^ Since then, the expression of c-*fos* and other immediate early genes (IEGs) has become a widely used surrogate for neuronal activity, transforming both molecular neuroscience and circuit behavioral studies. IEG-based methods robustly report neuronal activation, however, decreases in activity are not always translated into diminished IEG signals, making them less ideal for detecting the inhibition of neuronal activity (broadly defined here as the decrease of firing from baseline). To date, there has been no report of an equivalent indicator for detecting a decrease of neuronal activity (i.e., inhibition) as a trackable molecular event.

To fill this long-standing gap and advance our understanding of the inhibitory side of bi-directional neuronal activity dynamics, we sought to identify an endogenous molecular event associated with the inhibition of neuronal activity through unbiased screens. An ideal marker should be temporally linked to the decreased firing of action potentials yet result in molecular changes that are amenable to stable and trackable detection. The slower kinetics of *de novo* gene expression events makes them less suitable for this application. By contrast, protein post-translational modifications (PTMs), such as phosphorylation, are known to be robustly regulated by cellular activities in a faster and bi-directional manner. Therefore, we focused our search on activity regulated protein phosphorylations through quantitative TMT (Tandem Mass Tag) proteomics. Through a set of optogenetically enabled phosphoproteomic screens, we found that phosphorylation of pyruvate dehydrogenase E1 subunit alpha 1 (or pPDH) inversely correlates with the intensity of action potential firing in neurons both in cultured neurons and *in vivo*. We demonstrate that pPDH reliably detected neuronal inhibition in a range of contexts, thus establishing pPDH as the first trackable, endogenous, molecular event representing neuronal inhibition.

## Results

### Enable an optogenetic-based quantitative phosphoproteomic screen

To identify a molecular marker of inhibition, we performed an unbiased phosphoproteomic screen to search for endogenous cellular events induced by inhibition. Because robust TMT-proteomics requires a large amount of material, traditionally, proteomic profiling of neuronal activity is achieved by KCl depolarization or electrical stimulation, neither of which is ideal for bi-directional activity modulation. Such modulation can be achieved, however, by optogenetics, which enables precise, fast control of circuit activities in live animals. We therefore sought to adapt and scale optogenetics in cultured neurons to generate synchronized, precise firing patterns in a large enough number of neurons to provide sufficient materials for TMT-based phosphoproteomic profiling. We designed an optogenetic platform in which primary cortical neurons were transduced with light-activated ChR2, the most used channelrhodopsin (Figure 1A). We quantified the firing of action potentials (APs) under different light conditions by electrophysiological recordings. We found that 10 ms, 5 mW/mm^2^ 488 nm light stimulation over a 10 cm^2^ area reliably and precisely elicited APs up to 10 Hz^2^ (Figure 1B-C and S1A), achieving synchronized APs in >10^6^ neurons at the same time. As a positive control, we confirmed that such optical stimulation induced robust mRNA expression of well-established IEGs, such as *c-fos, Arc*, and *Npas4* after one hour (Figure 1D).

**Figure 1.**
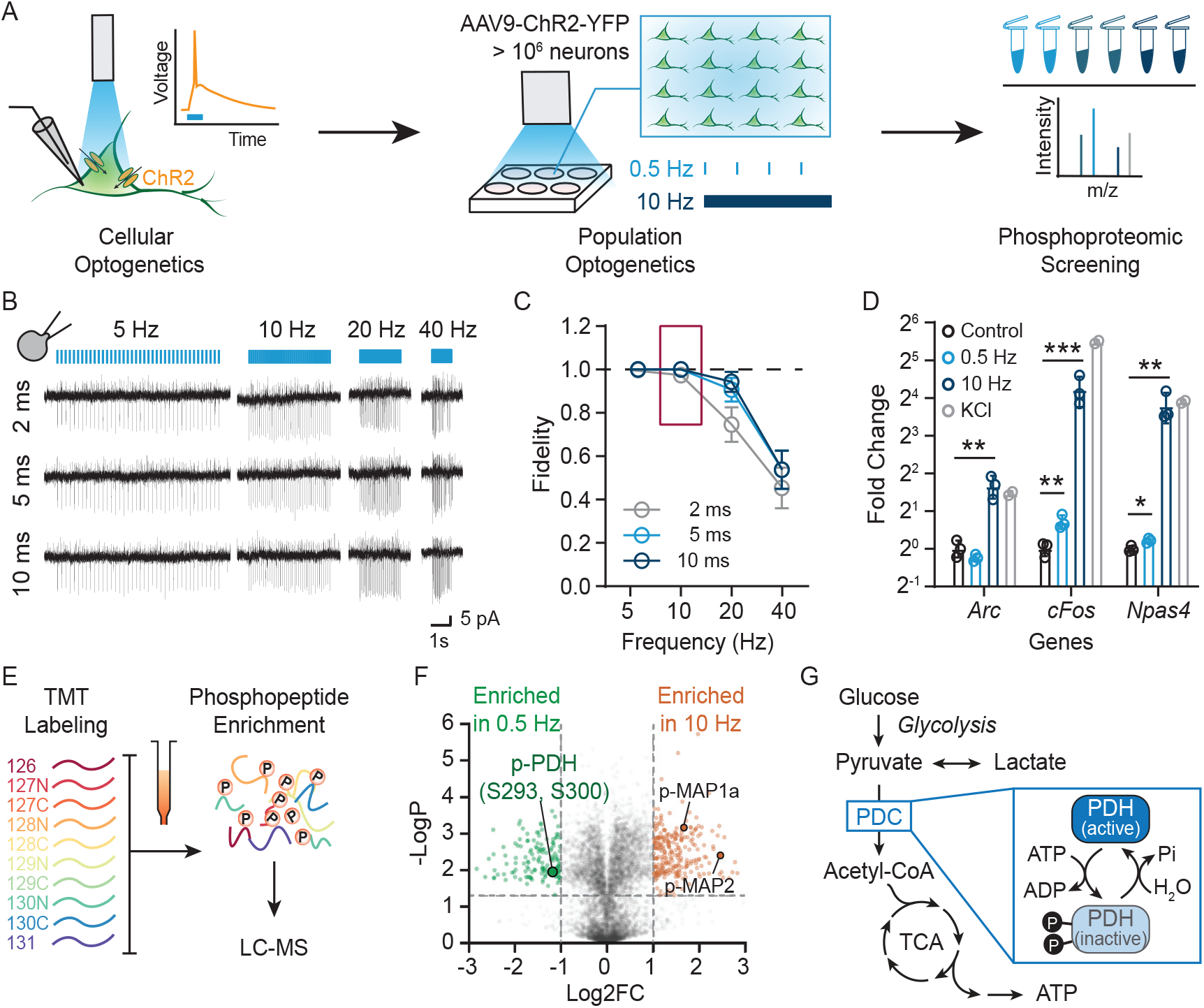
Activity-dependent phosphoproteomic screen identifies phospho-PDH in vitro. (See also Figure S1) (A) Schematic of the activity-dependent phosphoproteomic screen. Cultured cortical neurons infected with AAV9-Channelrhodopsin 2 (ChR2) were stimulated with blue light at different frequencies. Cells were harvested after stimulation for biochemical analysis. (B-C) Validation of optical spiking in cultured cortical neurons expressing ChR2. (B) Representative light-evoked traces using cell-attached recording at the indicated light pulse frequencies and durations (triggered by 40 pulses, λ = 470 nm, 5 mW/mm^2^). (C) Quantification of light-evoked spike fidelity in (B). N = 21 neurons per condition. (D) Gene expression changes of immediate early genes after light stimulation in cultured cortical neurons expressing ChR2. (N = 3 for control, 0.5 Hz, 10 Hz, N = 2 for KCl). (E) Workflow of phosphoproteomic analysis. Cells stimulated with different frequencies were lysed followed by tandem mass tag (TMT)-based labeling. TMT-labeled peptides were then enriched for phosphopeptides. (F) TMT-based phosphopeptide quantification revealed pPDH was significantly enriched in the 0.5 Hz stimulated group compared to 10 Hz stimulation group. (7,504 plotted phosphopeptides, cutoff fold change = 2, *p* = 0.05). (G) Schematic overview of glucose metabolism regulated by pPDH. All values are mean ± s.e.m. Statistics determined by two-tailed unpaired t test in (D). * p < 0.05; ** p < 0.01; *** p < 0.001.

### Phosphorylation of PDH is negatively correlated with neuronal activity

Initially, we set up an optogenetic paradigm to model slow (0.5 Hz) and fast (10 Hz) AP firing, followed by TMT-based untargeted phosphoproteomics for differentiating phosphopeptide changes associated with these activity patterns. We identified ∼7,543 phosphopeptides (Figure 1E and 1F). We detected ∼50 significantly changed phosphorylation targets, including MAP1a (Microtubule Associated Protein 1A) and MAP2, both of which have been reported to have activity-regulated phosphorylation^3^ (Figure 1F). We also identified two phosphorylation sites (Ser293 and Ser300) on Pyruvate Dehydrogenase E1 Subunit Alpha 1a (PDHA1a), which we refer to as pPDH^4^, that were more abundant in the low activity group compared to the high activity group (Figure 1F).

PDHA1a is the rate-limiting enzyme of the pyruvate dehydrogenase complex (PDC), which controls the entry of pyruvate into the oxidative TCA cycle (Figure 1G). Neuronal firing uses a large amount of energy, and it has been demonstrated that this energy is mostly provided by the TCA cycle. Although the coupling mechanism is still not understood, it is clear that activation of TCA is required for neuronal firing^5,6^. Phosphorylation of PDH inhibits enzymatic activity^7^, whereas its dephosphorylation activates PDH, subsequently increasing the production of ATP (Figure 1G). This is consistent with our finding that pPDH was reduced significantly when neurons were firing at a high rate (Figure S1B). We hypothesize that upon high neuronal activity, PDH is dephosphorylated to ramp up ATP production to meet the increased energy demand. By contrast, inhibition quickly leads to the re-phosphorylation of PDH as the energy demand drops to keep the baseline energy consumption low.

### pPDH specifically and dynamically correlated with neuronal activity in primary neurons

To test this hypothesis beyond the initial high throughput screens, we next sought specifically examine the relationship between PDH and neuronal activity in the primary neuron culture system. Both phosphorylated S293 (pS293) and S300 (pS300), two well-characterized phosphorylation sites of PDH, were detected in primary neurons by proteomics. However, upon examination of the tissue lysate from mouse prefrontal cortex, visual cortex, hippocampus, and cerebellum, pS293 appears to be the predominate form *in vivo* (Figure S2E and S2F), hence we decided to focus on pS293 PDH. Although in the initial screens we used 10 Hz optogenetic stimulation to maximize the difference between groups, here we adapted to a milder photostimulation protocol (5 Hz) to better resolve the quantitative dynamics of pPDH. Consistent with our proteomic findings, we observed that neuronal activation (5 Hz) leads to a decrease in pPDH in primary neurons by western blot (Figure S2A and S2B). We verified the specificity of antibody detection using a well-established PDK inhibitor, dichloroacetate(DCA), which blocks pPDH, and by using multiple antibodies (Figure S2C and S2D)^7^.

Biochemical assays using cultured neurons allow reliable and quantitative determination of the dynamic relation between neuronal activity and pPDH. Optogenetic- and KCl-induced neuronal activation led to a significantly lower pPDH level compared to stimuli of other signaling pathways (Figure 2A-C), suggesting reduced pPDH is specific to neuronal activity. We further characterized the temporal dynamics of activity-induced pPDH changes. Under 5 Hz continuous optogenetic stimulation conditions, we observed decreased pPDH in less than 10 minutes (Figure 2D-F). 50 minutes after the cessation of activation, pPDH level returned to a level comparable to the baseline (Figure 2G-I).

**Figure 2.**
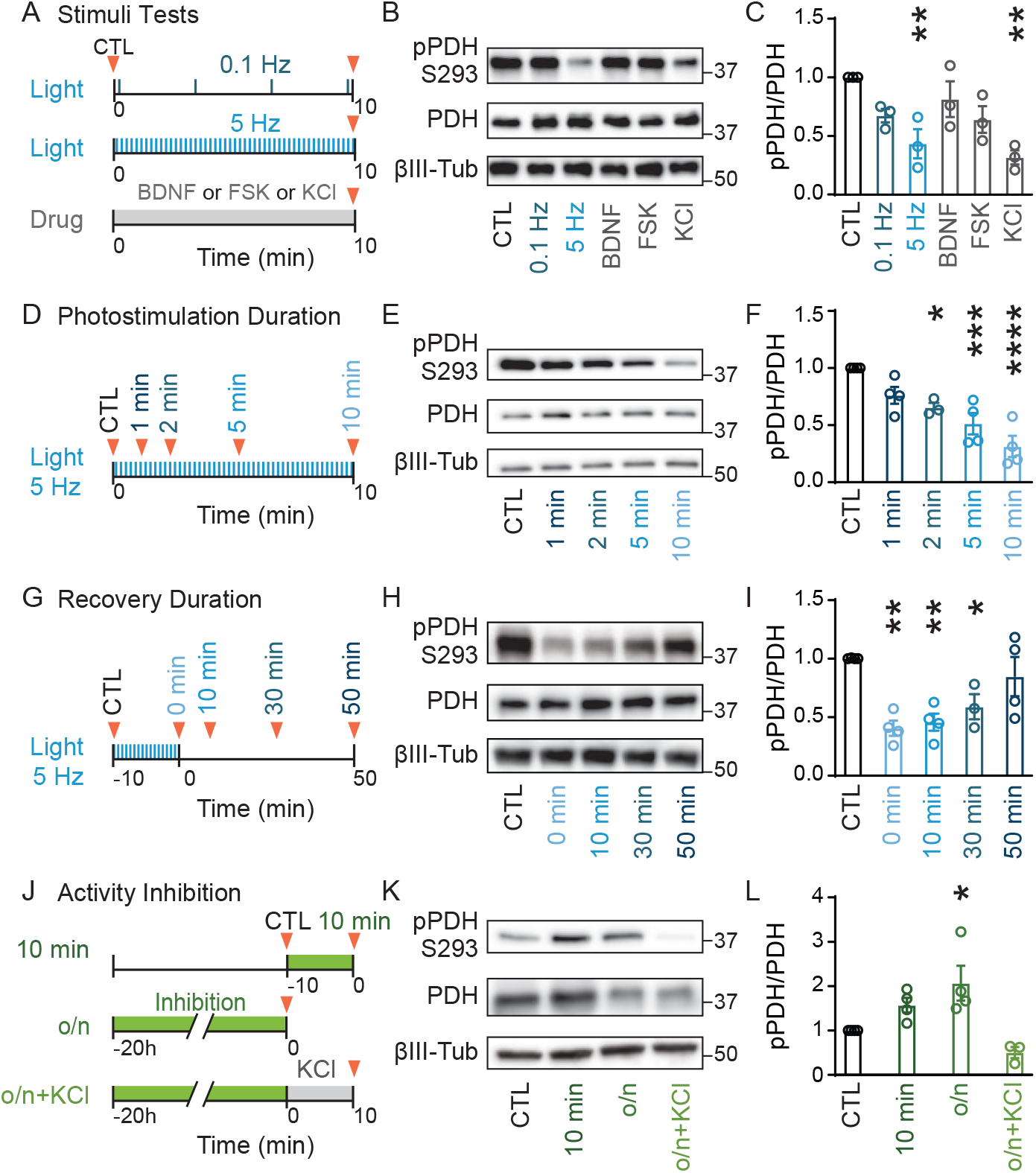
pPDH is inversely correlated with neuronal activity in vitro. (See also Figure S2) (A-C) The effects of different cellular stimuli on pPDH. (A) Cultured cortical neurons were applied with either light (0.1/5 Hz) or BDNF (50 ng/mL), Forskolin (FSK,10 μM) or KCl (50 mM) for 10 minutes. Cells were lysed at the end of stimulation (marked by red triangles) for immunoblot analysis (B), and pPDH levels were quantified in (C). N = 3 replicates per condition. (D-F) The effect of photostimulation duration on pPDH. (D) Cultured cortical neurons were stimulated with light (5Hz) for indicated durations. Whole-cell lysates were analyzed by immunoblotting (E), and pPDH levels were quantified in (F). N = 3 for 2 minute group and N = 4 for other groups. (G-I) The effect of post-stimulation recovery time on pPDH. Cultured cortical neurons stimulated with light (5Hz, 10 minutes) were allowed to recover for the indicated time (G) before whole-cell lysis and immunoblot analysis (H) pPDH levels were quantified in (I). N = 3 for 30 minute group and N = 4 for other groups. (J-L) The effect of pharmacological inhibition on pPDH. (J) Cultured cortical neurons were treated with an inhibitor cocktail (50 μM AP5, 10 μM CNQX, and 1 μM TTX) for 10 minutes or overnight (o/n). Whole-cell lysates were analyzed by immunoblotting (K), and pPDH levels were quantified in (L). N = 3 for o/n + KCl (50 mM) and N = 4 for other groups. All values are mean ± s.e.m. Statistics determined by ordinary one-way ANOVA with Dunnett’s multiple comparisons test. * p < 0.05; ** p < 0.01; *** p < 0.001; **** p < 0.0001, between the control and each treatment group. BDNF: brain-derived neurotrophic factor; FSK: forskolin.

To directly test how the decrease of activity affects pPDH, we took advantage of the high spontaneous activity in cultured primary neurons. A pharmacological inhibitory cocktail (10 uM CNQX, 50 uM AP5, and 1 uM TTX) was used to suppress spontaneous activity in primary neurons. Acute (10 minute) and chronic (overnight) pharmacological inhibition led to an increase in pPDH, which can be reversed by the addition of KCl (Figure 2J-L). Together, these results demonstrate that pPDH can be bidirectionally and inversely regulated by neuronal activity.

### Immunofluorescence labeling of pPDH marks neuronal inhibition

After biochemically establishing the correlation between pPDH and neuronal activity in cultured primary neurons, we next tested if pPDH can serve as a histological label for neuronal inhibition *in vivo*. We first examined pPDH levels in response to general anesthesia by isoflurane, which globally inhibits brain activity. As previously reported, 2 hours of general anesthesia led to widespread loss of cFOS immunofluorescence (IF) across brain regions^8,9^. We detected a robust increase of pPDH IF in many cortical and subcortical brain regions (Figure 3A-C and S3A).

**Figure 3.**
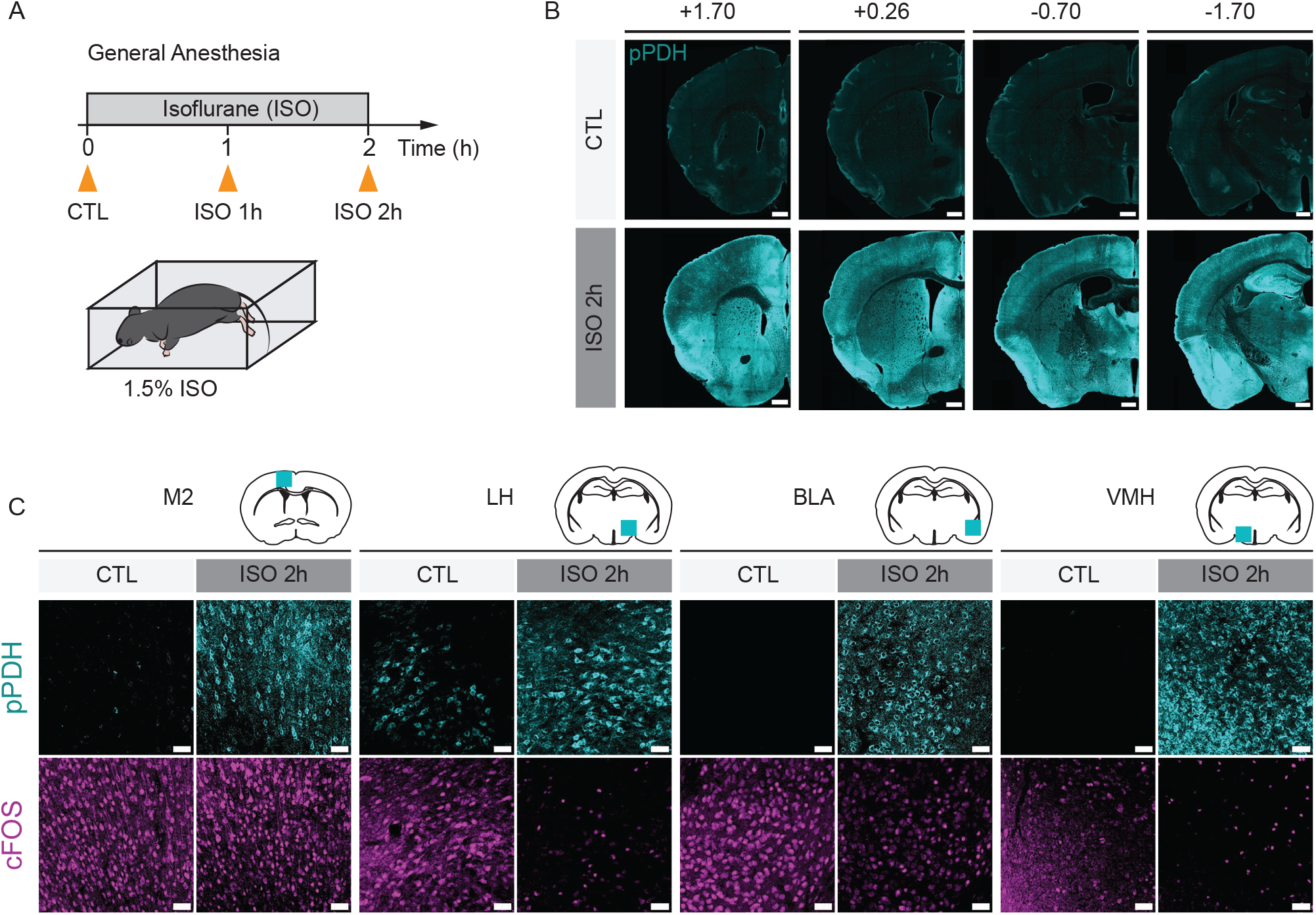
pPDH inversely correlates with neural activity in general anesthesia. (See also Figure S3) (A) General anesthesia paradigm. Mice were exposed to isoflurane, and brain samples were collected at indicated time points (orange triangles) for immunohistochemistry analysis. (B-C) pPDH labeling in the whole brain after 2 hours of general anesthesia. (B) Representative images of brain coronal sections at indicated bregma positions. (C) Representative zoomed-in images of pPDH and cFOS labeling in different brain regions. N = 4 animals per condition. Scale bar: 500 μm in (B) and 50 μm in (C). M2: secondary motor cortex; LH: lateral hypothalamic area; BLA: basolateral amygdaloid nucleus, anterior part; VMH: ventromedial hypothalamic nucleus.

Consistent with an earlier report^8^, we also observed that general anesthesia led to a higher cFOS signal in the supraoptic nucleus (SO), where high pPDH was detected in non-overlapping neurons in the same area (Fig S3B). Note not all regions showed higher pPDH labeling, for example, the CA3 region of the hippocampus did not display a significant change in pPDH level despite showing decreased cFOS signal under general anesthesia (Fig S3C), suggesting that similar to IEGs, pPDH labeling may have regional preference and variation. Interestingly, at an earlier time point (1 hour after anesthesia), strong pPDH was already detected across the brain, but the loss of cFOS was less obvious (Figure S3D and S3E), reflecting the faster dynamic nature of pPDH compared to IEG expression.

In addition to general anesthesia, we wanted to test how pPDH alters in response to more specific neuronal activity changes within a circuit and when induced by controlled sensory experiences. To this end, we adapted a well-established visual stimulation protocol and examined pPDH dynamics in the mouse visual circuits^10,11^. Mice were first kept in the dark for 48 hours (visual deprivation) and then exposed to bright visual stimulation for 4 hours to induce neuronal activity, before going back to the dark again for 2 hours and 12 hours, at which point the visual stimulation-induced activity is expected to subside (Figure 4A). Consistent with the literature^10,11^, the cFOS signal in the primary visual cortex (V1), lateral geniculate nucleus (LGN), and lateral entorhinal cortex (LEnt) was robustly induced by the 4-hour visual stimulation and slowly returned to baseline only after 12 hours in the second dark phase. By contrast, the pPDH signal in all three regions showed a sharp increase 2 hours after the mice were re-introduced to the dark, suggesting that cessation of visual stimulation (and hence decreased activity in the visual pathway) can be rapidly captured by pPDH (Figure 4B-C and S4).

**Figure 4.**
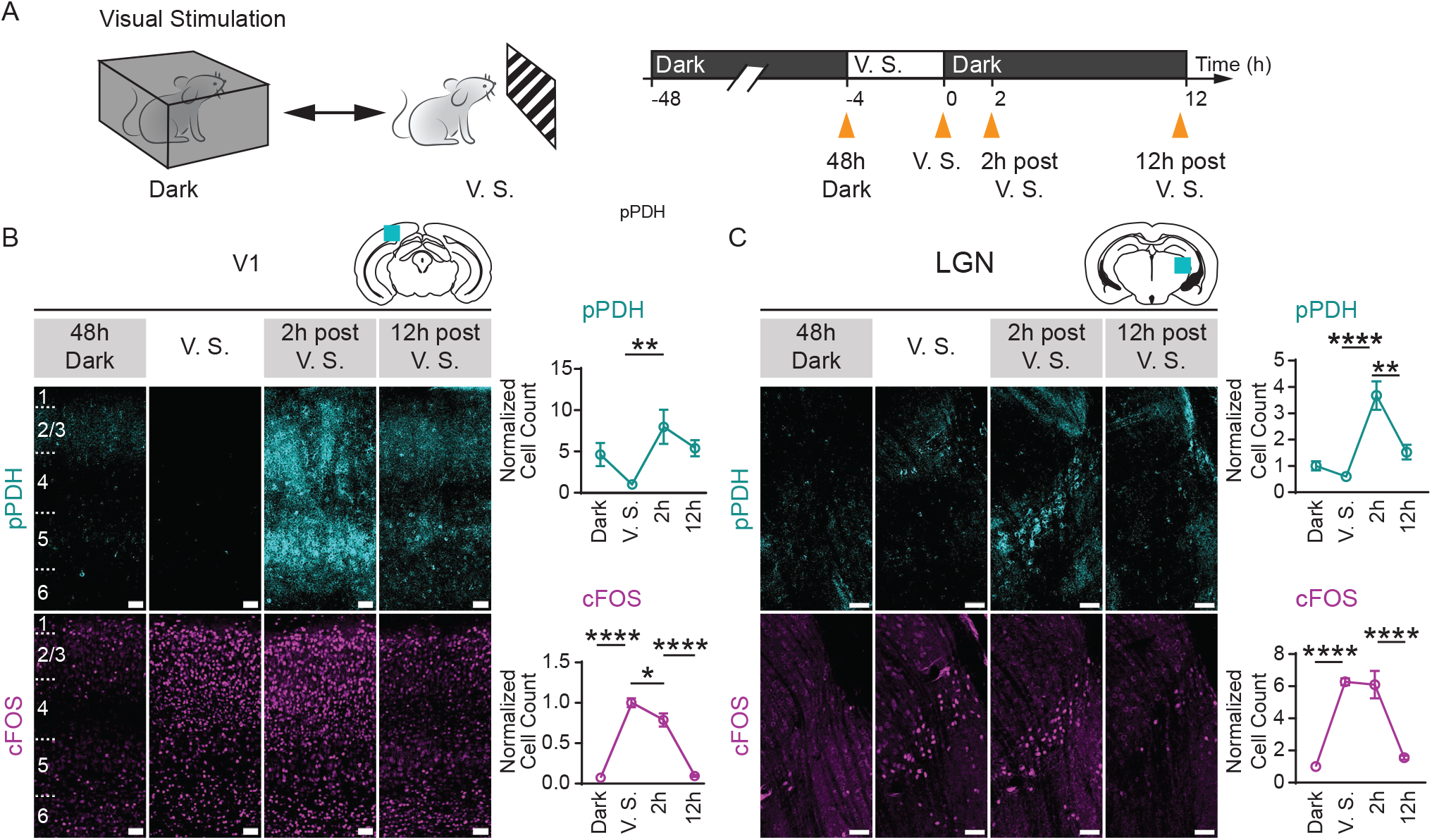
pPDH inversely correlates with neural activity in visual stimulation. (See also Figure S4) (A) Visual stimulation paradigm. Mice were kept in the dark for 48 hours before being exposed to visual stimuli (V. S.) for 4 hours and then returned to the dark immediately thereafter. Tissues were harvested at different timepoints (orange triangles) for immunostaining. (B) Representative images and quantification of pPDH and cFOS staining in V1. Positively stained cells in the layer 5 of V1 were quantified. The cell counts were normalized to the V.S. group. 48 hour dark (N = 6), V. S. (N = 10), 2 hours post V.S. (N = 7), 12 hours post V.S. (N = 6). (C) Representative images and quantification of pPDH and cFOS staining in LGN. The cell counts were normalized to the 48 hour dark group. N=4 for each group. All values are mean ± s.e.m. Statistics determined by one-way ANOVA with Tukey’s multiple comparisons test. * p < 0.05; ** p < 0.01; *** p < 0.001; **** p < 0.0001. Scale bar: 50 μm in (B-C). V1: primary visual cortex; LGN: lateral geniculate nucleus.

### pPDH marks *in vivo* neuronal inhibition by natural behaviors and internal states

Beyond sensory-induced activity, we next examined if behavior or internal state-induced activity and inhibition can be represented by pPDH. We first examined the median preoptic nucleus (MnPO), a key region for water homeostasis, using a well-established water deprivation and drinking model^12,13^. Mice were water deprived for 24 hours before regaining access to water (Figure 5A). As reported, MnPO activity was induced by water deprivation, as demonstrated by robust cFOS labeling in this region. It is also known that MnPO activity is quickly suppressed upon drinking^12,13^, which was rapidly captured by increased pPDH staining 1 hour after water access. By contrast, cFOS signal slowly returned to baseline 8 hours after drinking, again reflecting the difference in the off-kinetics between pPDH and cFOS labeling (Figure 5A).

**Figure 5.**
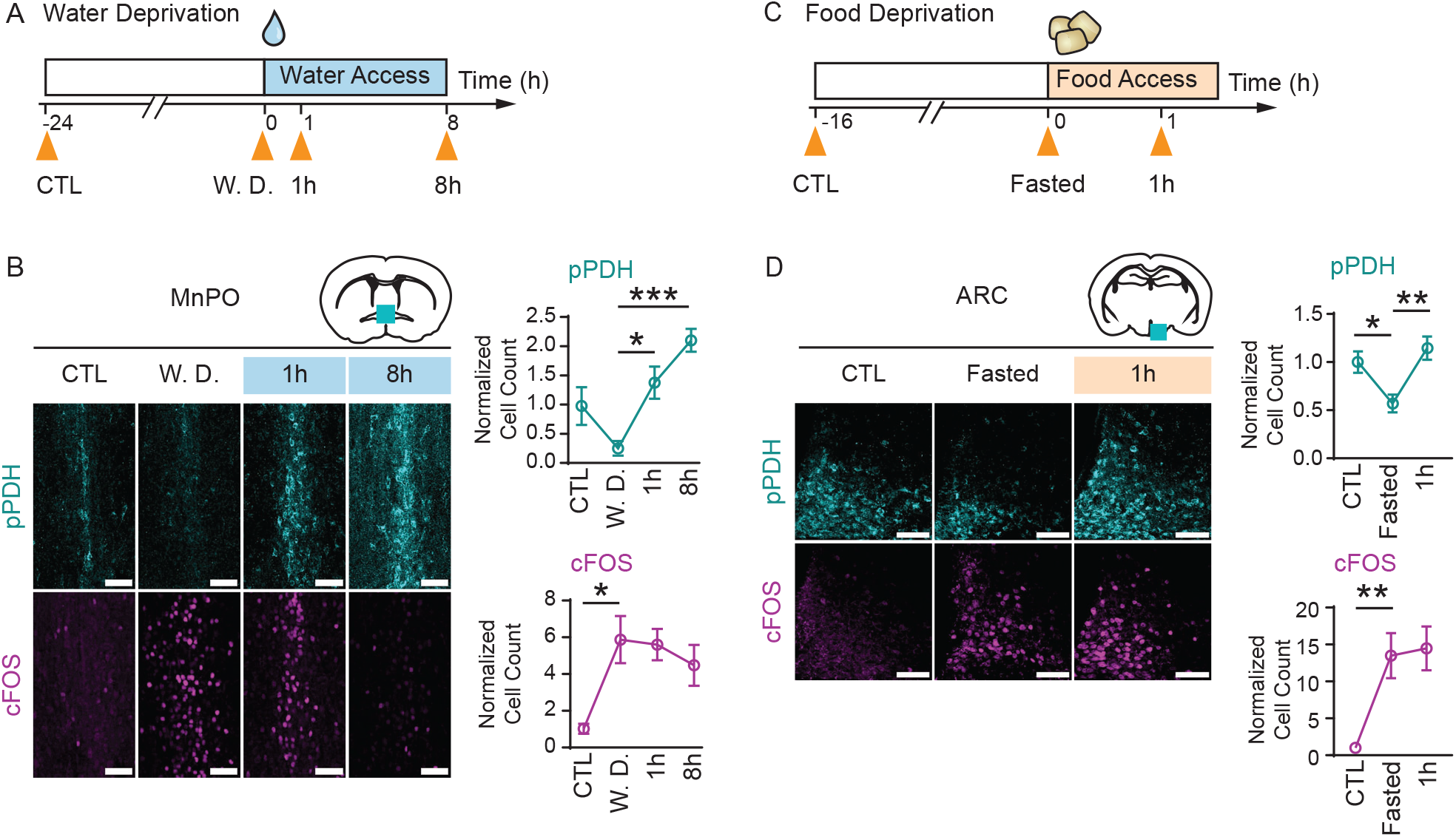
pPDH inversely correlates with neural activity in water or food deprivation models. (See also Figure S5) (A-B) Water deprivation paradigm. (A) Mice were water deprived for 24 hours before given access to water. Brain samples were collected at indicated timepoints (orange triangles) for immunohistochemistry analysis. (B) Representative images of pPDH and cFOS staining in MnPO. Positively stained cells were quantified and normalized to the control group. N = 4 animals per group. (C-D) Fasting-refeeding paradigm. (C) Mice were food deprived for 16 hours before given access to food for 1 hour. Brain samples were collected at indicated timepoints (orange triangle) for immunohistochemistry analysis. (D) Representative images of pPDH and cFOS staining in ARC. Positively stained cells were quantified and normalized to the control group. N = 10 for control groups and N = 11 for fasted and 1 hour refed groups. All values are mean ± s.e.m. in (B) and (D). Statistics determined by ordinary one-way ANOVA with Tukey’s multiple comparisons test. * p < 0.05; ** p < 0.01; *** p < 0.001. Scale bar, 50 μm. MnPO: median preoptic nucleus; ARC: arcuate hypothalamic nucleus.

Another robust model of internal state-induced behavior is fasting and re-feeding. Overnight fasting induces strong activation of the agouti-related protein (AgRP) neurons in the arcuate hypothalamic nucleus (ARC), as confirmed by the robust increase in cFOS signal in the ARC (Fig 5B). Conversely, we observed a decrease in pPDH signal in the same region after fasting. Similar to the MnPO neurons, refeeding rapidly suppresses AgRP neuron activity^14,15^. Although this is well documented by *in vivo* calcium imaging experiments (Figure S5A)^14,16^, we could not detect significant decreases in cFOS signal in the ARC 1 hour after refeeding (Figure 5B), suggesting that it likely takes much longer for the cFOS protein to return to baseline. By contrast, strong pPDH induction is apparent in the ARC 1 hour after refeeding. Using AgRP-Cre mice, we further demonstrated that these pPDH+ ARC neurons are mostly AgRP+ neurons (Figure S5B), fitting their calcium dynamics as detected by *in vivo* fiber photometry. Together, these data show that pPDH provides a way to histologically label neuronal inhibition previously inaccessible by existing IEG-based activity markers and directly demonstrate the feasibility of using pPDH to track the reduction of neuronal activity *in vivo*.

### pPDH reveals cell-type neural dynamics associated with feeding behaviors

A key advantage of histologically compatible markers is that they offer an unbiased way to identify new neural substrates associated with behaviors or experiences. For example, c-*fos* and other IEGs have proven to be powerful tools for illuminating how different brain regions and neuron types respond to various stimuli, behavior, or experiences. We therefore tested the ability of pPDH to reveal dynamics of novel brain regions or cell types.

In our fasting-refeeding experiments designed to characterize the ARC-AgRP neurons, we noticed that a group of discrete neurons displayed significant and specific pPDH decrease with fasting but quickly rebounded upon refeeding in the LH (Figure 6A and S6A), suggesting they were activated by fasting but inhibited by refeeding. However, such inhibition was not detected by cFos immunostaining (Figure 6A) or other IEG-based methods in the literature. We took advantage of that pPDH staining can be readily multiplexed with other histological labels to identify the cell types in this region. Upon dual-labeling the refeeding induced pPDH+ cells with three well-characterized LH cell-type markers including orexin (hypocretin), melanin-concentrating hormone (MCH), and vesicular GABA transporter (Vgat), we found that the pPDH+ neurons highly overlapped with orexin (137/164 pPDH+ cells) but not MCH (3/132) or Vgat (5/169) neurons (Figure 6B). These results were consistent with a previous *in vivo* calcium imaging report on the inactivation of orexin/hypocretin neurons during eating^17^. Thus, using pPDH as an inhibition marker, we were able to find a population of orexin/hypocretin LH cells that are activated by fasting but rapidly inhibited by refeeding (Figure S6B).

**Figure 6.**
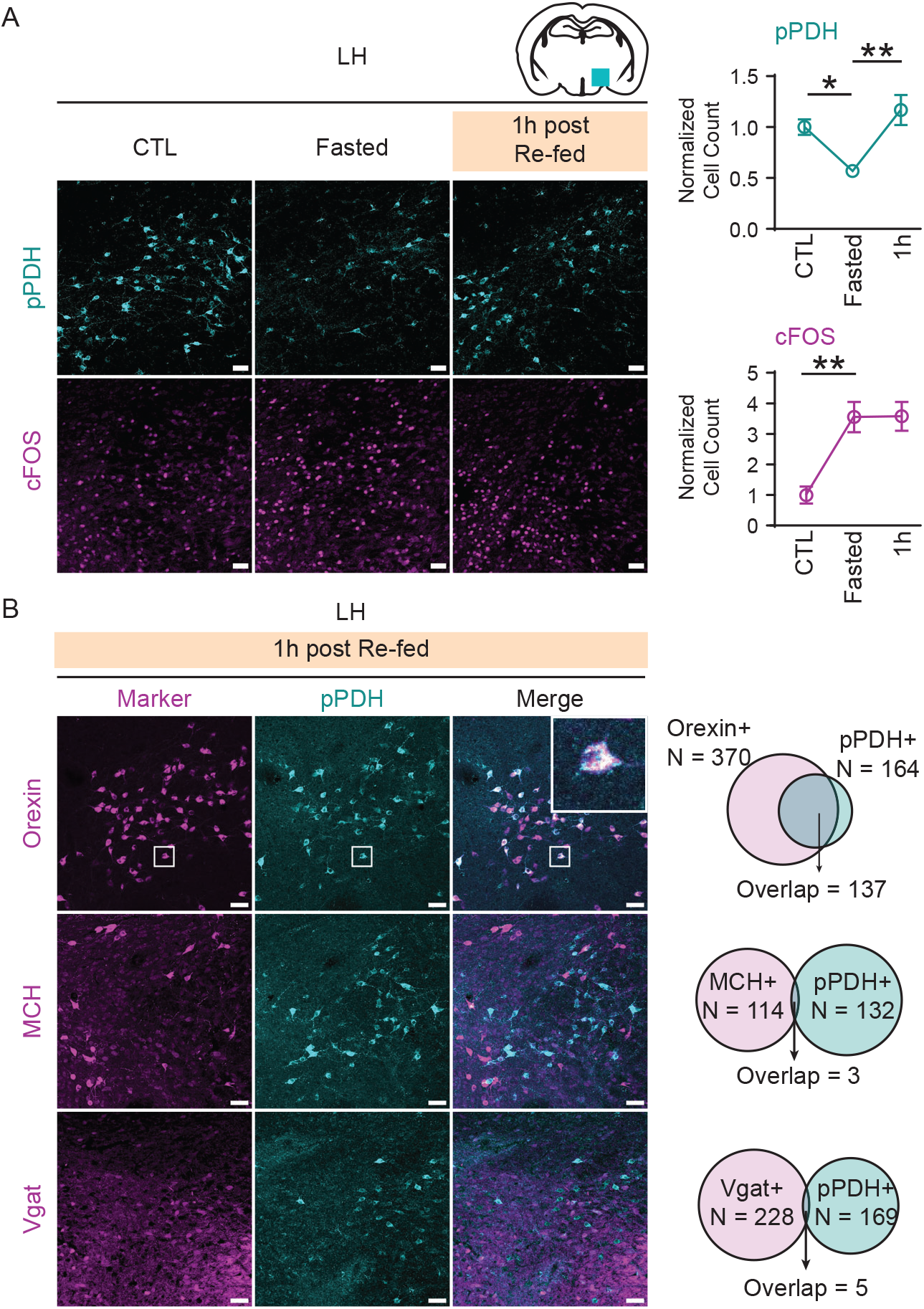
pPDH inversely correlates with neural activity in LH in fasting-refeeding paradigm. (See also Figure S6) (A) Representative images of pPDH and cFOS staining in LH. Positively stained cells were quantified and normalized to the control group. N = 4 animals per group. (B) Representative images and quantification of pPDH labeling and cell-type markers in LH 1 hour after refeeding. Zoomed-in views of co-labeled cells are shown in insets. N = 3 mice for Orexin and MCH (by orexin and MCH immunostaining), N = 4 mice for Vgat (using Vgat-Cre: Ai9 mice). All values are mean ± s.e.m. in (A). Statistics determined by ordinary one-way ANOVA with Tukey’s multiple comparisons test. * p < 0.05; ** p < 0.01. Scale bar, 50 μm in (A) and (B). LH: lateral hypothalamic area.

## Discussion

Neuronal dynamics involve both the increase and decrease of activities, yet for a long time, there have only been trackable histological markers for the former. Indeed, the existing literature is heavily centered on the activation of neural dynamics, because in general, it has been difficult to screen and track the reduction of neuronal activity with traditional markers. This imbalance highlights the need for novel tools to profile, track and study this critical part of circuit dynamics. To this end, we searched for a reverse marker of neuronal activity, focusing on PTMs, which are modulated on a time scale closer to the inhibition of neuronal activity. By using an optogenetic approach, we were able to perform phosphoproteomics on differentially activated neurons, leading to the identification of pPDH as a marker inversely correlated with neuronal activity.

The directionality and dynamics of pPDH fit the energetic requirements for neural firing— dephosphorylation activates PDH, which gates pyruvate entry to the TCA cycle for full oxidation and ATP production, to meet the high demand for energy when activity is high. Our findings suggest that PDH is rapidly re-phosphorylated when activity ceases or decreases, presumably to avoid wasting energy. This dynamic regulation of neuronal metabolism gave us a molecular handle to probe and track the firing state of the neurons.

Although we identified pPDH as an inverse activity marker in an unbiased way, there is long-standing evidence of its involvement in neuronal activity. As early as 1982, a highly abundant ∼40kD phosphoprotein from brain lysate, later determined to be pPDH, was found to be visible by eye on protein gel and influenced by electrical stimulations^18^. This underscores that pPDH is not simply a bystander molecular event, but a prominent and logical link between biochemical changes and electrical activity.

Our findings have multiple important implications. First, by establishing an optogenetics-mediated proteomic screen, for the first time, we enabled genetically defined and optically controlled high-throughput profiling of neuronal activities. This approach could be subsequently expanded to cell-type-specific and activity pattern-specific optogenetic screens with numerous applications. Second, through the identification of pPDH, we now have a trackable endogenous marker to *post-hoc* identify inhibited neurons. Although inhibition can be readily read out from recording or Ca^2+^ imaging, it remains difficult to track and isolate those inhibited neurons for subsequent molecular, genetic, or projection studies; unlike activated neurons, which can often be “trapped” by c-*fos* or other IEG derivatives. Now with pPDH, it is feasible to conduct inhibition profiling at the same scale and throughput as well as with the same cell identification compatibility as can be achieved with IEG staining. For example, combining pPDH staining with whole brain clearing, imaging, or screening tools would transform our ability to understand bi-directional changes of activity, study neuronal populations with increased refinement, and unravel circuits underlying both well-established and new paradigms.

### Limitations of the study

First, we detected pPDH using a monoclonal antibody, which works robustly with conventional histology or can be combined with newer spatial imaging techniques, especially when combined with dual-color IEG staining in the same sample. However, compared to the highly developed IEG toolbox (including but not limited to Fos-EGFP^19^, Fos-LacZ^20^, Fos-ChR2^21^, Arc-Venus^22^, E-SARE^23^, TRAP^10^/TRAP2^24^, CANE^25^, Fos-tTA^26^, CAPTURE^27^, RAM^28^), pPDH-based detection is still at its infancy and currently limited to post-mortem brains. Nonetheless, we expect additional protein and genetic engineering will render new capabilities to pPDH-based indicators, similar to the development and refinement of IEG tools in the last few decades. Second, like any indicators, we expect that the baseline and sensitivity of pPDH could vary across different brain regions and cell types. While it is not feasible to exhaustively test all potential biases in this study, we did notice, for example, that the baseline pPDH level was high in the hippocampus and not responsive to isoflurane (Figure S3B), limiting the use of pPDH as an indicator in this region. Thus, it is important to consider potential regional preference as a caveat in future pPDH applications.

## Acknowledgments

We thank all members of the Ye lab and the Dorris Neuroscience Center for their support and feedback. We thank Min Huang and Kathryn Spencer for their technical support. L.Y. is supported by the National Institutes of Health Director’s New Innovator Award (DP2DK128800), NIDDK (DK114165 and DK124731), the Dana Foundation, the Whitehall Foundation, and the Baxter Foundation. D.Y., Y.W., Z.P., N.K.L. were supported by the Dorris Scholar Award.

## Author contributions

LY and YW conceived the study. DY, YW, TQ and LY designed the study. DY, YW, TQ, XZ, LS, JM, ZP, NKL, DBM, KW, YX, FP, AM, VA, HTC, and JRY performed the experiments and analyzed the data. XZ, DBM, and JRY performed proteomics studies. Funding acquisition: LY. Projection administration: LY. Supervision: LY, JRY, HTC, VA, AM. DY, YW, and LY wrote the manuscript with input from all authors. All authors reviewed and provided feedback on the manuscript.

## Lead contact

Further information and requests for resources and reagents should be directed to and will be fulfilled by the Lead Contact, Li Ye.

## Materials availability

This study did not generate new unique reagents.

### Data and code availability

The original phosphoproteomic data prior to analysis and the mass spectrometry data reported in this paper will be deposited to MassIVE upon acceptance.

## EXPERIMENTAL MODEL AND SUBJECT DETAILS

### Primary cortical neuron culture

Primary cultured cortical neurons were prepared from P0 C57BL/6J mouse pups of both sexes. Cortexes were isolated, digested with papain (2.5 mg/mL, Worthington) supplemented with DNase I (0.5 mg/mL, Millipore Sigma), and plated onto glass coverslips (Neuvitro) or 35 mm dishes (Corning) precoated with poly-D-lysine (0.1 mg/mL, Millipore Sigma). Cultures were maintained at 37°C and 5% CO2 with growth medium (Neurobasal-A media (Gibco) containing GlutaMAX (1%, Gibco) and B-27 supplement (2%, Gibco)). Fluorodeoxyuridine (FUDR, 20 μM, Millipore Sigma) was added into the medium at DIV3 after plating to inhibit growth of astrocytes in neuronal cultures. The neurons were then transduced with AAV (AAV9-hSyn-hChR2(H134R)-EYFP) in growth medium with a constant multiplicity of infection [MOI (defined here as vector genomes per cell)] of 10,000 vector genomes/cell to achieve at least 70% transfection rate based on EYFP expression prior to in vitro population optogenetic stimuli and electrophysiology experiments. Electrophysiology recording and population optogenetics were performed between DIV14-16.

### Animals

All animal experiments were performed in accordance with the National Institutes of Health Guide for the Care and Use of Laboratory Animals and approved by Scripps Research Institutional Animal Care and Use Committee (IACUC 18-0001). Unless specified, adult mice of both genders (6-12 weeks of age) were used for in vivo studies. C57BL/6J (Jackson Laboratory; C57BL/6J; stock #000664) mice were used for general anesthesia, visual stimuli, water deprivation and drinking, and fasting-refeeding experiments.

AgRP-Cre (Jackson Laboratory; Agrptm1(cre)Lowl/J; stock #012899) mice were used for fiber photometry recording in food deprivation. Ai9 mice (Jackson Laboratory; B6.Cg-Gt(ROSA)26Sortm9(CAG-tdTomato)Hze/J; stock #007909) were crossed with AgRP-Cre or Vgat-Cre (Jackson Laboratory; Slc32a1tm2(cre)Lowl/J; stock #016962) for cell identity mapping in food deprivation experiments. Mice were group housed and maintained on a 12h light: 12h dark cycle with food and water ad libitum unless specified. Mice were randomly assigned to experimental groups.

## METHOD DETAILS

### *In vitro* population optogenetic stimulation

Primary neurons were prepared and transduced as described above. Inhibitor cocktail containing 6-cyano-7-nitroquinoxaline-2,3-dione (CNQX; Tocris, 10 μM), (2R)-2-amino-5-phosphonopentanoic acid (AP5; Abcam, 50 μM) and Picrotoxin (Abcam, 100 μM) in culture media was applied the day before population optogenetic stimulation. An LED light (Prizmatrix) mounted on a homemade holder was used to deliver even light (λ = 455 nm) to a square surface to cover one 35-mm dish. The LED was connected to a Master-8 or a Digidata 1440A digitizer to generate pulses at desired frequency. Light pulses (10 ms, 5 mM/mm^2^) were delivered at indicated frequency and duration for phosphoproteomic screening and western blot experiments. After stimulation, the neurons were immediately processed to RNA extraction, phosphoproteomic screen, or western blot.

### *In vitro* pharmacological treatment

Primary neurons were prepared and transduced as described above. For brain derived neurotrophic factor (BDNF), forskolin (FSK), KCl or dichloroacetate (DCA) treatment, the neurons were treated with 10 μM CNQX, 50 μM AP5 and 100 μM Picrotoxin overnight before pharmacological treatment. BDNF (50 ng/mL, Pepro Tech), FSK (10 μM, Sigma), KCl (50 mM, Sigma) or DCA (stock solution 12M, added to indicated concentration) was added to culture media as previously described^29^ and left for 10 min before the neurons were harvested. For activity inhibitor experiment, the neurons were treated with 50 μM AP5, 10 μM CNQX, and 1 μM tetrodotoxin (TTX) in culture media for overnight or 10 min. After pharmacological treatment, the neurons were immediately processed to RNA extraction, phosphoproteomic screen, or western blot.

### Primary culture electrophysiology

Primary neurons were prepared and transduced as described above, and treated with 10 μM CNQX, 50 μM AP5 and 100 μM Picrotoxin overnight. Recording of neurons prepared and transduced as above were obtained in Tyrode’s solution (in mM: 140 NaCl, 5 KCl, 2 CaCl_2_, 0.8 MgCl_2_, 10 HEPES and 10 Glucose, pH 7.4, 300 mOsm) supplemented with 10 μM CNQX, 50 μM AP5 and 100 μM picrotoxin, with an internal solution (in mM: 120 K-gluconate, 20 KCl, 4 Na_2_ATP, 0.3 Na_2_GTP, Na_2_ phosphocreatine, 0.1 EGTA and 10 HEPES, pH 7.3, 305 mOsm) in 3-6 MΩ glass pipettes. ChR2-expressing neurons were visually identified by expression of EYFP. Light from a 470 nm LED (Thorlabs) with light power density adjusted of 5 mM/ mm^2^ were used for blue light illumination to evoke action potentials. All data collected from cell-attached recordings. Recordings were made using Multiclamp 700B amplifier (Molecular Devices). Signals were filtered at 3kHz and sampled at 5kHz with a Digidata 1440A analog-digital interface (Molecular Devices). pClamp 10.7 software (Molecular Devices) was used to record and analyze data.

### Phospho-proteomic profiling of cultured neurons

#### Sample preparation

Primary neurons were prepared and transduced as above. After in vitro population optogenetic stimulation, cells were washed with ice-cold PBS and scraped into lysis buffer (1 mM EPPS, 8 M urea, protease (complete mini, EDTA-free) inhibitors (Roche) and PhosSTOP phosphatase inhibitor (Roche)). Cells were further probe-sonicated (6 mA for 2-3 sec, ∼10 rounds) on ice. Lysate concentrations were measured with BCA protein assay and adjusted to 1 mg/mL. Per 100 ug lysate, the proteins were reduced with 10mM tris(2-carboxyethyl)-phosphine (TCEP) (Sigma) for 20 min, alkylated with 10 Mm iodoacetamide in dark for 20 min, and then quenched with 10 mM dithiothreitol (DTT) (Sigma) for 20 min on a shaker at room temperature (RT). Lysates were sequentially mixed with 4 volumes of MeOH, 1 volume of CHCl_3_ and 3 volumes of H_2_O. The mixture was vortexed and centrifuged to afford a protein disc at the interface of CHCl_3_ and H_2_O. The protein disc was recovered, washed with ice-cold methanol and air dried. The resulting protein pellets were resuspended in EPPS buffer (100 μL, 10 mM, pH 8), digested with Lys-C (Wako Chemicals, 1:50 (protease to protein)) overnight, then with trypsin (Promega, 1:50 (protease to protein)) for 6 hours on a 37 °C shaker.

#### TMT labeling and peptide fractionation

Digested peptides (5 conditions, 4 biological replicates/condition) were split into two groups, each containing 2 replicates for each condition. Each group of peptides were labelled with 10 TMT reagents from a 11-plex reagent kit (Thermo A34808 and A34807) as directed by the manufacturer. Each 0.8 mg vial of TMT reagent was dissolved with 44 μL anhydrous acetonitrile, equivalent to four aliquots of 0.2 mg. Digests were each reacted with a channel of TMT label in final 30% (v/v) acetonitrile at RT for 1 hour (0.1 mg peptides per 0.2 mg TMT label). Labeling was quenched with 0.3% (w/v) NH_2_OH at RT for 15-20 min, and then acidified with 3% formic acid to pH ∼2.5. For a 11-plex experiment. Samples (10%/channel) were combined as the Total (T) sample and fractionated into 10 parts using high pH reversed phase peptide fractionation kit (Thermo 84868) as directed by the manufacturer.

#### Phosphopeptide enrichment

The remaining TMT-labeled peptides were desalted by passing through Sep-Pak C18 cartridges (Waters). The desalted peptides enriched for phosphopeptides using Hi Select Fe-NTA Phosphopeptide enrichment kits (Thermo A32992) as directed by the manufacturer. Briefly, the peptides were loaded to Fe-NTA spin column, washed and eluted. The eluent was evaporated to rdyness using SpeedVac vacuum concentrator, and redissolved in buffer A (95% H_2_O, 5% CH_3_CN, 0.1% FA), and passed through Pierce C18 Spin Tips (Thermo 87782). The flow through from Fe-NTA columns was subjected to a second enrichment using Hi Select TiO_2_ Phosphopeptide enrichment kits (Thermo A32993) following manufacturer’s protocol.

The eluted phosphopeptides were combined with the previous enriched peptides, and fractionated into 7 parts using a high pH reversed phase peptide fractionation kit eluted with a series of 20 mM NH_4_HCO_3_ / CH_3_CN buffers. The solvent of each fraction was removed using SpeedVac vacuum concentrator. The resulting fraction was redissolved in buffer A and ready for LC-MS analysis.

#### Liquid chromatography and mass spectrometry (LC-MS)

Samples were analyzed by liquid chromatography tandem mass-spectrometry using an Orbitrap Fusion mass spectrometer (Thermo Scientific) coupled to an EASY-nLC 1200 UPLC autosampler chromatography (Thermo Scientific). The peptides were loaded directly to a C18 analytical column (100 um × 25 cm, tip diameter 5 um) packed with BEH 1.7 um beads (Waters), and separated using buffer A and buffer B gradients (buffer A: 95% H_2_O, 5% CH_3_CN, 0.1% FA; buffer B: 20% H_2_O, 80% CH_3_CN, 0.1% FA). TMT-labelled phosphopeptides used the following gradient: 5-40% buffer B in buffer A from 0-100 min, 40-90% buffer B from 100-130 min, 90% buffer B from 130-140 min. T sample peptides used the following gradient: 1-30% buffer B in buffer A from 0-160 min, 30-90% buffer B from 160-220 min, 90% buffer B from 220-240 min. The voltage applied to nanoelectron spray ionization (nESI) source was 2.8 kV and the source temperature was 275 °C. Peptide spectra were acquired using the data-dependent acquisition (DDA) synchronous precursor selection (SPS)-MS3 method. Briefly, the scane began with an MS1 scan (Orbitrap analysis, resolution 120000, automatic gain control AGC target 4e5, max injection time 50 ms, m/z 400-1500). The most intense precursor ions at charge state 2-7 were isolated by the quadrupole and CID MS/MS spectra were acquired in the ion trap in Turbo scan mode (isolation width 1.6 Th, CID collision energy 35%, activation Q 0.25, AGC target 1e4, maximum injection time 100 ms, dynamic exclusion duration 10 s), and finally 10 notches of MS/MS ions were simultaneously isolated by the orbitrap for SPS HCD MS3 fragmentation and measured in the orbitrap (60k resolution, isolation width 2 Th, HCD collision energy 65%, m/z 120-500, maximum injection time 120 ms, AGC target 1e5, activation Q 0.25). Quality control LC-MS/MS of digests before TMT labeling was acquired on an Orbitrap Velos mass spectrometer coupled with an EASY-nLC 1200 UPLC autosampler chromatography.

#### MS data processing and analysis

Spectra were analyzed using the Integrated Proteomics Pipeline (IP2) platform. MS data were searched using the ProLuCID algorithm against a sequence database of UniProt SwissProt *Mus musculus* reviewed proteome appended with the sequences of common contaminant proteins, and with the reverse sequences as decoy (UniProt downloaded 2018-11-28, total 34189 protein entries). TMT-peptide CID MS/MS spectra were searched and filtered using the following parameters: static modifications of TMT (229.1629, N-term, Lys) and carbamidomethylation (57.02146, Cys), dynamic oxidation on Met (15.9949), phosphorylation on S/T/Y (79.9663), precursor mass tolerance of 10 ppm, fragment ion mass tolerance 0.6, one or no tryptic end (K/R), up to 3 missed cleavages, and minimum peptide length 4.

Protein identification required at least 1 peptide identified per protein, and was filtered to peptide and protein false discovery rate < 1% using the target-decoy method.

Filtered phosphopeptides were quantified in IP2 based on the MS3 TMT reporter ion intensity. To identify upregulated/downregulated phosphopeptides between two conditions (i.e. 10 Hz vs 0.5 Hz, or 10 Hz vs control), two-tailed t tests (two-sample unequal variance) were performed on the two conditions dataset. Phosphopeptides with statistically significant changes were identified by Fold change > 2 (upregulated) or < 0.5 (downregulated) and p < 0.02.

### RNA preparation and RT–qPCR analysis

Primary neurons prepared and transduced as above were stimulated by in vitro population optogenetics. After stimulation, neurons were washed in ice-cold PBS and proceeded to total RNA extraction using RNeasy Mini kits (Qiagen). Total RNA was reverse-transcribed using High Capacity cDNA reverse transcription kit (Applied Biosystems #4368813). The resultant cDNA was mixed with primers (Integrated DNA Technology) and SyGreen Blue Mix (Genesee Scientific, 17-507) for RT–qPCR using the CFX384 real-time PCR system (BioRad). Normalized mRNA expression was calculated using ΔΔC_t_ method, using Tbp (encoding TATA-box-binding protein) mRNA as the reference gene. Statistical analysis was performed on ΔΔC_t_.

qPCR primers used (5’ to 3’): *Tbp* (CCTTGTACCCTTCACCAATGAC and ACAGCCAAGATTCACGGTAGA); *Fos* (GGGAGGACCTTACCTGTTCG and AGGCCAGATGTGGATGCTT); *Npas4* (CAGATCAACGCCGAGATTCG and CACCCTTGCGAGTGTAGATGC).

### Western blot analysis

Primary neurons prepared and transduced as above were stimulated by in vitro population optogenetics or pharmacological treatment. After stimulation, neurons were washed in ice-cold PBS, lysed in RIPA buffer (G-Biosciences) supplemented with 1x Halt protease inhibitor cocktail (Thermo Scientific) and 1x Halt phosphatase inhibitor cocktail (Thermo Scientific) by vortex and sonication. For mouse brain tissue samples, the brains were dissected, washed in ice-cold PBS and cut into 1-mm thick coronal sections. The selected brain regions were harvested using a 2-mm diameter tissue punch, and homogenized in RIPA buffer (1x Halte protease inhibitor cocktail, 1x Halt phosphatase inhibitor cocktail). Protein concentrations were quantified using BCA assay (Thermo Scientific).

Each sample contains 10-15 ug of total protein, 4x Laemmli loading buffer (Bio-Rad) was added, and the samples were heated at 75 °C for 10 min. The protein samples were resolved using SDS-PAGE (10% acrylamide gel, Bio-Rad) and transferred to 0.45 um PVDF membranes (Millipore Sigma). The membrane was blocked with 5% BSA in Tris-buffered saline (TBS, Genesee Scientific) with 0.1% Tween 20 (TBST) buffer at RT for 1h, incubated with primary antibodies diluted in 5% BSA/TBST at 4°C overnight. The membrane was washed with TBST (3 times), then incubated with secondary antibody in 5% BSA/TBST at RT for 90 min. The membrane was then washed in TBST (3 times) and developed with ECL western blot substrate (Bio-Rad) and imaged using Azure C300 imaging system (Azure biosystems). Relative band intensities were quantified using ImageJ.

Primary antibodies used in this study include: rabbit polyclonal anti-PDH (1:1000; Proteintech #18068-1-AP) used in Figure 2 and S2D-E, rabbit polyclonal anti-pPDH (Ser293) (1:1000; Millipore #AP1062) in Figure 2, S2B and S2D-F, rabbit polyclonal anti-pPDH (Ser300) (1:1000; Millipore #AP1064) in Figure S2B and S2D-E, rabbit monoclonal anti-pPDH (Ser293) (1:1000; Cell Signaling #37115) in Figure S2F, rabbit polyclonal anti-beta III Tubulin (1:10000; Abcam #ab18207) in Figure 2, S2B and S2D-E. HRP-conjugated secondary antibodies (Jackson ImmunoResearch) were used at 1:10000.

### Behavior assays

All behavioral experiments were performed according to well-established protocols from the literature. Mice were subjected to only the minimal handling required for genotyping and colony maintenance prior to performing every experiments.

#### General anesthesia assays

Mice (male) were placed in the chamber on a 37°C heating pad with standard isoflurane vaporizer. Isoflurane (4% for induction (∼5 min), 1.5% for maintenance) was applied into the chamber^30^. After 1h or 2h of isoflurane exposure, mice were transcardially perfused with ice-cold PBS following ice-cold 4% paraformaldehyde (PFA). The dissected brains were further post-fixed for immunohistochemistry.

#### Visual deprivation and stimulation assays

Visual deprivation and stimulation assays were adapted from published protocol^10^. Briefly, the mice were group housed in transparent cages (Innovive) with free access to food and water and transferred into a dark room at 6 pm. The cages were then placed in a light-tight cubicle with complete darkness for 48h.

Light stimuli were delivered by 7-inch tablets mounted on all four sides of the cages, each of the tablets produces light of ∼385 lux. After 4h of visual stimuli, the mice were returned to complete darkness for 2 h or 12 h. At the indicated time points, mice were transcardially perfused and the dissected brains were proceeded for immunohistochemistry.

#### Water deprivation and drinking assays

Water deprivation and drinking assays were performed as previously described^12^. Briefly, mice were water deprived for 24h before giving free access to water. At the indicated timepoints, mice were transcardially perfused and the dissected brains were proceeded for immunohistochemistry.

#### Fasting-refeeding assays

Fasting-refeeding assays were adapted from previous report^31^. Mice were single housed 1 week before the experiment session. During the session, the mice were fasted overnight from 6pm to 10 am (ad libitum water). Mice were then given access to regular chow food. At the indicated timepoints, mice were transcardially perfused and the dissected brains were proceeded for immunohistochemistry.

### Immunohistochemistry

Mice were terminally anesthetized with isoflurane and intracardially perfused with PBS and 4% PFA. The brains were dissected and post-fixed in 4% PFA overnight at 4°C. 50 μm-thick coronal sections were prepared using a vibratome (Leica VT1000S) for immunohistochemistry. Free floating sections were blocked with 5% normal donkey serum in PBS with 0.3% Triton X-100 (PBST) at RT for 1h, and then incubated with primary antibodies diluted in PBST with 2% BSA at 4°C overnight. Slices were then washed by PBST for 3 times, and incubated with secondary antibodies (1:500, Jackson ImmunoResearch) diluted in PBST with 2% BSA at RT for 1h. After washing with PBST, slices were stained with DAPI (5nM in PBS) at RT for 15 min, and then mounted in fluoromount-G (Electron Microscopy Science) for confocal imaging.

For pPDH and cFos co-staining experiments, slices were stained with rabbit monoclonal anti-pPDH (Ser293, Cell Signaling #37115) and mouse monoclonal anti-cFos (Santa Cruz Biotechnology #sc-271243), washed in PBST, incubated with Alexa Fluor 488 conjugated donkey anti-mouse and Alexa Fluor 647 conjugated donkey anti-rabbit.

For pPDH and Orexin or MCH co-staining experiments, slices were firstly stained with rabbit anti-Orexin or anti-MCH, washed in PBST, then incubated with Alexa Fluor 488 conjugated donkey anti-rabbit secondary antibody. Slices were then blocked with 5% normal rabbit serum in PBST at RT for 1h, and then incubated with Alexa Fluor 647 direct-conjugated rabbit monoclonal anti-pPDH (Ser293) (#37115, custom conjugated by Cell Signaling).

The primary antibodies used include: rabbit monoclonal anti-pPDH (Ser293) (1:500, Cell Signaling #37115), mouse monoclonal anti-cFos (1:500, Santa Cruz Biotechnology #sc-271243), rabbit anti-Orexin (1:100, Cell Signaling #16743), rabbit anti-MCH (1:200, Phoenix Pharmaceuticals #H-070-47).

### Confocal microscopy

Mounted brain slices were imaged with Olympus FV3000 confocal microscope with ×10/0.6 NA water immersion objective (XLUMPlanFI, Olympus). Images were acquired using Fluoview.

### Stereotactic surgery and fiber photometry analysis

Induction chamber was used to anesthetize the AgRP-Cre mice using isoflurane. Mice were head fixed on a stereotactic device (Kopf) using ear bars. An incision was made on mice head to expose the skull. Using bregma and lambda system mice head was aligned. A small craniotomy was made above the site of injections (Bregma: −1.6 mm, midline: −0.25 mm, dorsal surface: −5.8 mm). 300 nl of AAV9-CAG-Flex-GCaMP6m-WPRE-SV40 (8×10^12^ GC per ml) virus was injected at 50 nl min^−1^ using a Nanoject II (Drummond) injector. Fiber photometry fiber optic cannula (400 μm diameter, 0.5 N.A., RWD life sciences) was implanted at (bregma: −1.6 mm, midline: −0.25 mm, dorsal surface: −5.7 mm) and secured to skull with dental cement. Mice were allowed to recover for 1-2 weeks before experiments or experiment-related habituation.

For recording AgRP activity during fasting-refeeding experiments, mice were fasted overnight for 16 hrs. Next day, mice were placed into a recording chamber with patch fiber (400 μm core NA 0.5, RWD) attached to their optic cannula. The optic fiber was connected to a 470 nm and 410 nm light source. Mice were allowed to habituate for 30 minutes before recording. Food was given 15 minutes after starting the recording. Fiber photometry and acquisition setup has been previously described^27^. Signal was expressed in delta F/F where F represents baseline fluorescence. Video recording of behavior was used to manually annotate and timestamp the start of feeding behavior.

## QUANTIFICATION AND STATISTICAL ANALYSIS

For the quantification of western blot, the relative band intensity was quantified by ImageJ^32^. Intensity ratio between pPDH (S293) and total PDH was calculated and plotted. For quantification of immunohistochemistry, images were analyzed by ImageJ. Each brain region was identified by DAPI stain and referencing to Allen Brain Atlas (https://mouse.brain-map.org/). The brightness and contrast were normalized for all images within each experiment. Staining-positive cells were segmented using thresholding, and then counted by analyzing particle function in ImageJ. The number of overlapping cells were counted manually.

Unless otherwise specified, statistical analysis was performed using Prism 9 (Graphpad). For RT-qPCR analysis, two-tailed unpaired t-test was used to examine the difference between groups. For phosphoproteomic analysis, two-tailed t-tests (two-sample unequal variance) were used to determine significantly enriched peptides. For western blot analysis, ordinary one-way ANOVA with Dunnett’s multiple comparisons was used to determine significant changes from the control group. For immunohistochemistry analysis, ordinary one-way ANOVA with Tukey’s multiple comparisons test was used to determine significant changes between the two groups. Numbers of biological replicates are indicated in figure legends. All data presented as mean ± s.e.m. Indications of significance level are as follows: * p < 0.05; ** p < 0.01; *** p < 0.001; **** p < 0.0001

---

**Figure S1.**
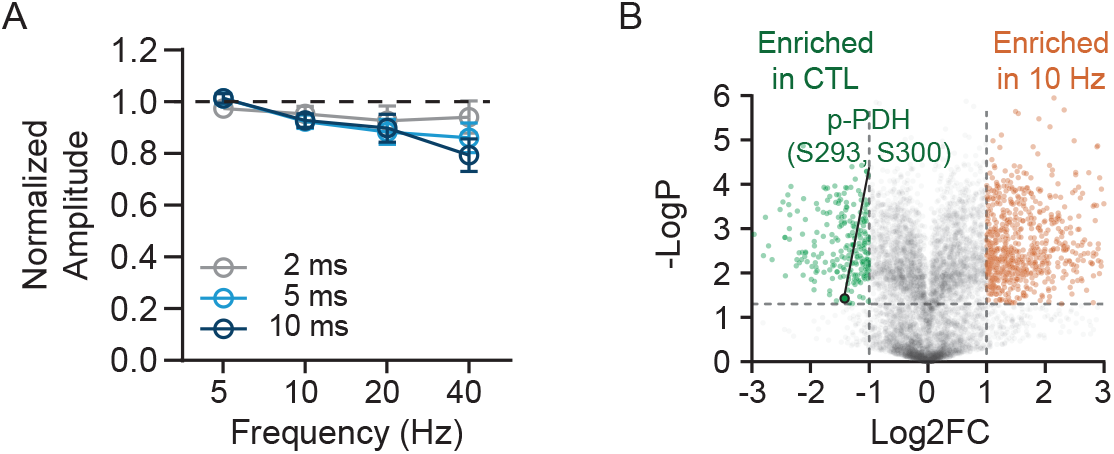
Additional analysis of activity-dependent phosphoproteomic screen, related to Figure 1. (A) Quantification of light-evoked spike amplitude using cell-attached recording in cortical neurons expressing ChR2. All values are mean ± s.e.m. (B) Volcano plot comparing phosphopeptides significantly changed in control and 10 Hz stimulated groups. pPDH was significantly enriched in the control group. (7,542 plotted phosphopeptides, cutoff fold change = 2, p = 0.05).

**Figure S2.**
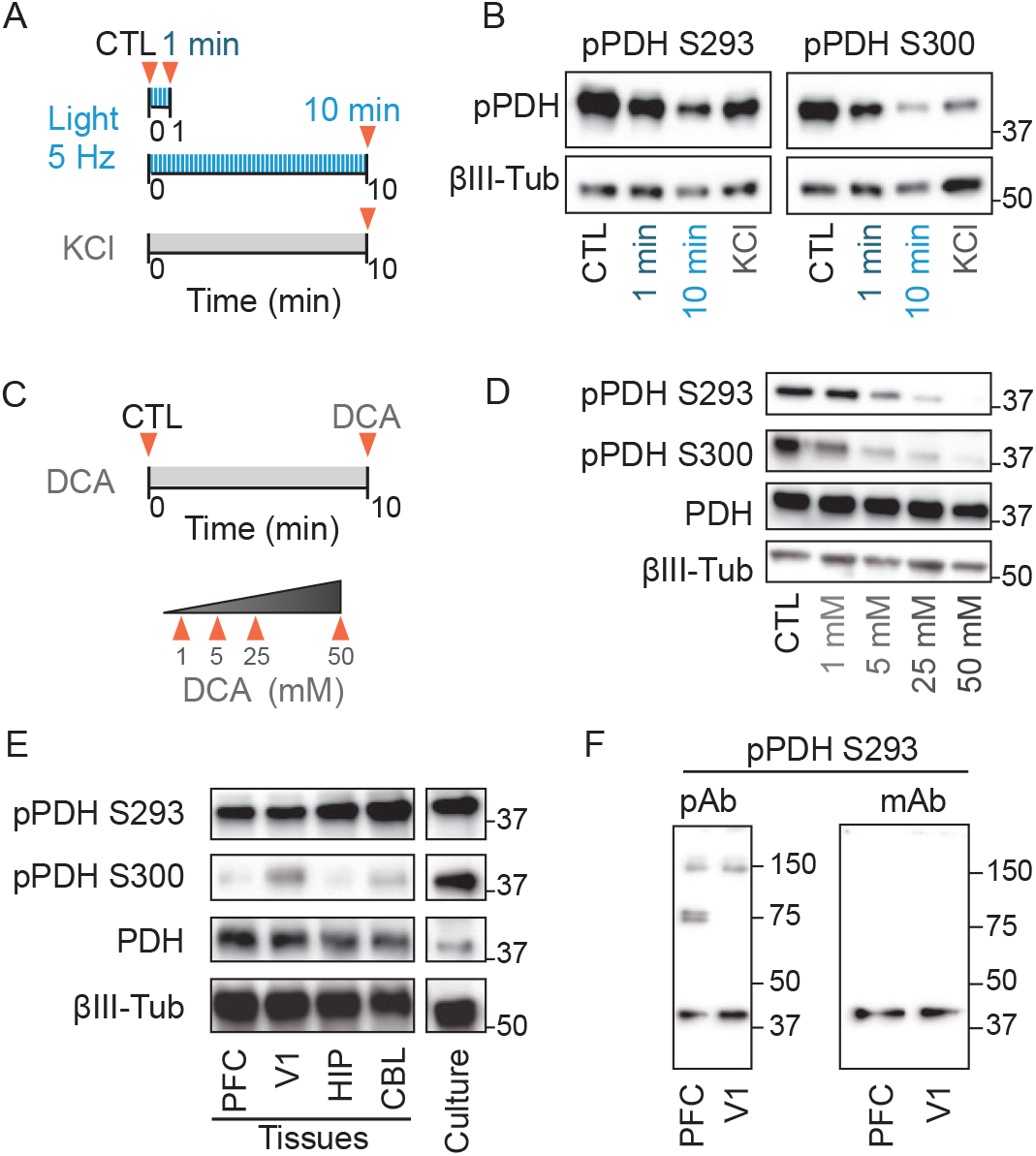
Additional characterization of stimulation effects on pPDH, related to Figure 2. (A-B) The effect of neuronal activity on pPDH. (A) Cultured cortical neurons were applied with either blue light (5 Hz) for 1 minute or 10 minutes, or KCl (50 mM) for 10 minutes. Cells were lysed at the end of stimulation (marked by red triangles) for immunoblot analysis (B). (C-D) The effect of pyruvate dehydrogenase kinase inhibitor (dichloroacetate (DCA)) on pPDH. (C) Cultured cortical neurons were treated with DCA at different doses, and cells were lysed for immunoblot analysis (D). (D) pPDH (S293) is the dominant phosphorylation site in vivo. Whole-cell lysates of mouse brain tissues (PFC, V1, HIP, and CBL) and cultured neurons were analyzed by immunoblotting for the phosphorylation sites of PDH. (E) Comparison of pPDH polyclonal (Millipore AP1062) and monoclonal (Cell Signaling 37115) antibodies by immunoblot analysis of PFC and V1 whole-cell lysates. PFC: prefrontal cortex; V1: primary visual cortex; HIP: hippocampus; CBL: cerebellum.

**Figure S3.**
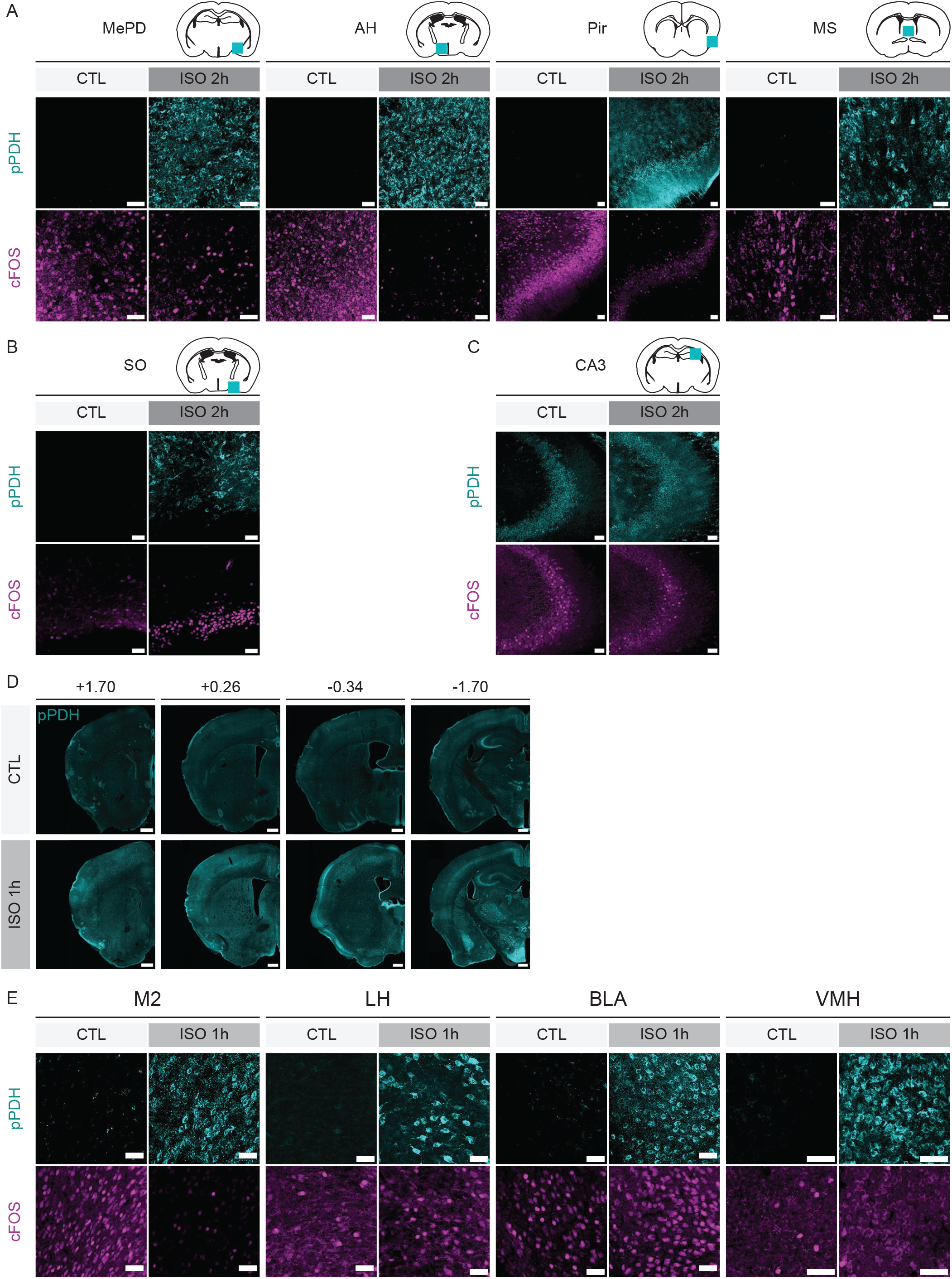
Additional characterization of pPDH in general anesthesia, related to Figure 3. (A-C) Representative zoomed-in images of pPDH and cFOS labeling in different brain regions after 2 hours of general anesthesia. (D-E) pPDH labeling in the whole brain after 1 hour of general anesthesia. (D) Representative images of coronal brain sections at indicated bregma positions after 1 hour of general anesthesia. (E) Representative zoomed-in images of pPDH and cFOS labeling in different brain regions. N = 3 mice for control and N = 4 mice for isoflurane treated group. Scale bar: 50 μm in (A-C and E), 500 μm in (D). MePD: medial amygdaloid nucleus, posterodorsal part; AH: Anterior hypothalamic area; Pir: piriform cortex; MS: medial septal nucleus; SO: supraoptic nucleus; CA3: field CA3 of hippocampus; M2: secondary motor cortex; LH: lateral hypothalamic area; BLA: basolateral amygdaloid nucleus, anterior part; VMH: ventromedial hypothalamic nucleus.

**Figure S4.**
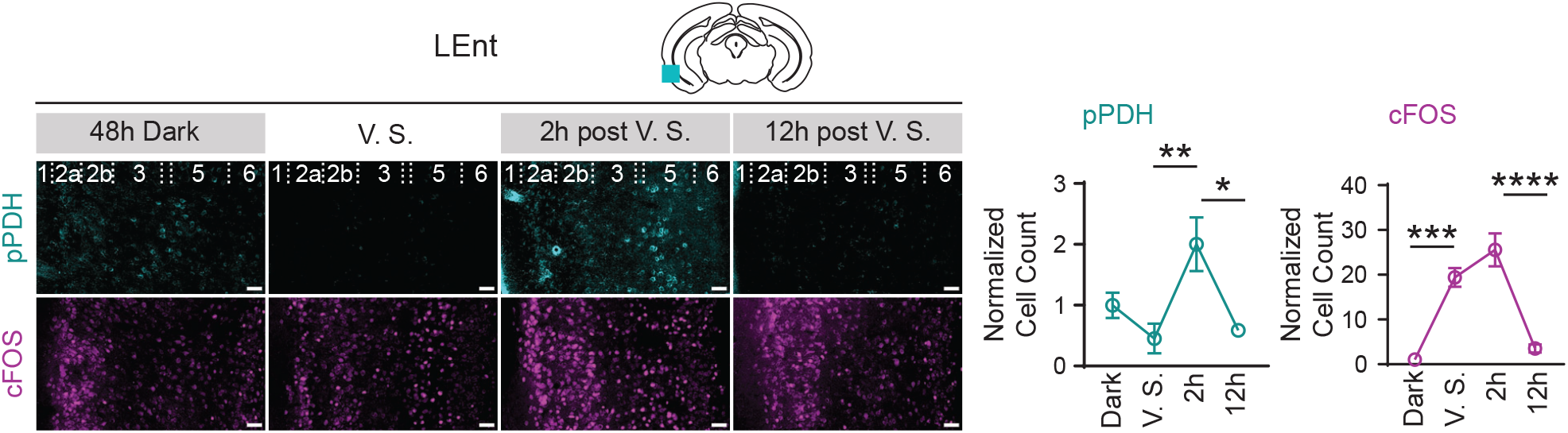
Additional characterization of pPDH in visual stimuli, related to Figure 4. Representative images of pPDH and cFOS staining in LEnt after visual stimuli. Positively stained cells in layer 5 were quantified and normalized to the 48 hour dark group. For LEnt: 48 hour dark (N = 4), V.S. (N = 4), 2 hour post V.S. (N = 4), 12 hour post V.S. (N = 4). All values are mean ± s.e.m. Statistics determined by ordinary one-way ANOVA with Tukey’s multiple comparisons test. * p < 0.05; ** p < 0.01; **** p < 0.0001. Scale bar: 50 μm. LEnt: lateral entorhinal cortex.

**Figure S5.**
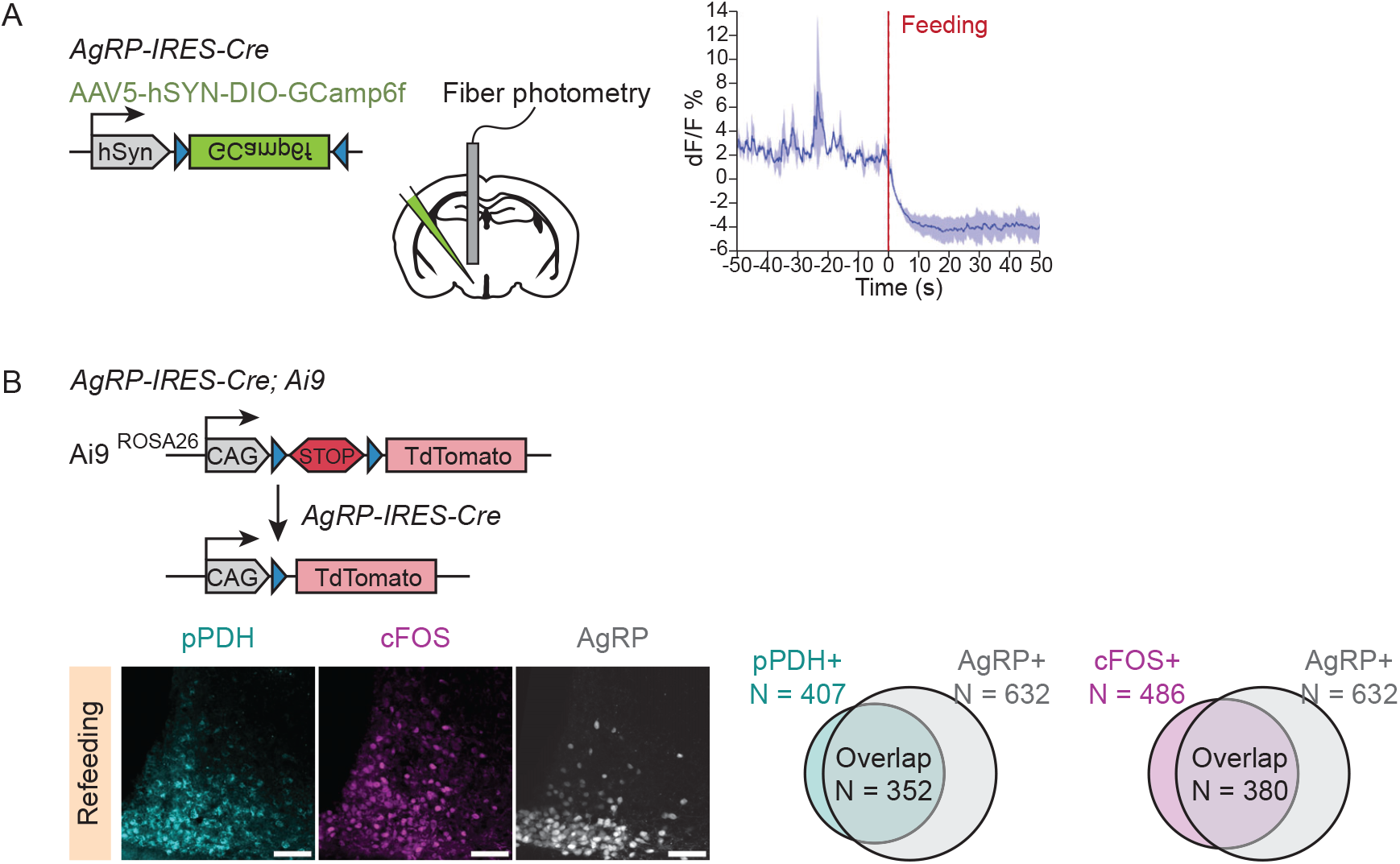
Additional characterization of pPDH labeling in the ARC, related to Figure 5. (A) Left, schematic of virus injection and fiber photometry. Right, Calcium signal from AgRP neurons in fasted mice presented with food. (N = 2 mice). (B) Top, schematic of the genetic strategy used to generate AgRP labeling mice. Bottom, representative images of pPDH and cFOS labeling in ARC of AgRP-Ai9 mice 1 hour after feeding. pPDH and cFOS labeled cells were quantified. N = 4 mice.

**Figure S6.**
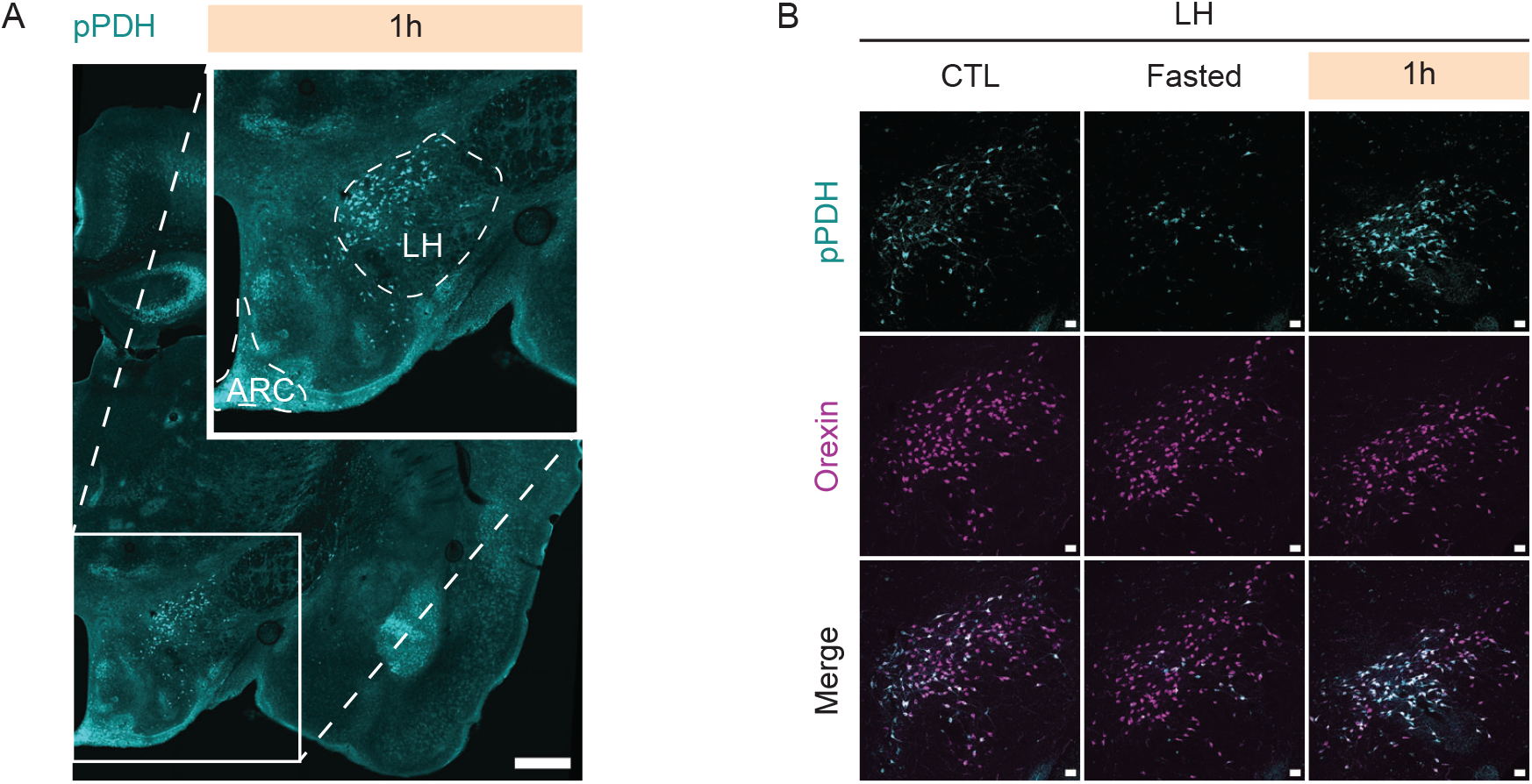
Additional characterization of pPDH labeling in the LH, related to Figure 6. (A) Representative image of brain coronal section showing pPDH labeling in ARC and LH in 1 hour refed mice. (B) Representative images of pPDH and Orexin labeling in LH in fasting-refeeding paradigm. Scale bar, 500 μm in (A), 50 μm in (B). ARC: arcuate hypothalamic nucleus; LH: lateral hypothalamic area

