## Supplementary_Figs for "Phosphorylation of pyruvate dehydrogenase marks the inhibition of *in vivo* neuronal activity"

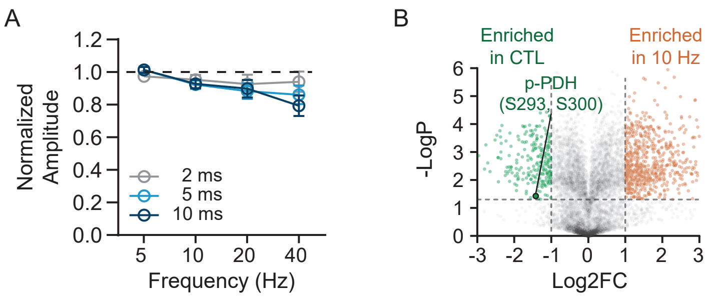


**Figure S1. Additional analysis of activity-dependent phosphoproteomic screen, related to Figure 1**

(A) Quantification of light-evoked spike amplitude using cell-attached recording in cortical neurons expressing ChR2. All values are mean ± s.e.m.

(B) Volcano plot comparing phosphopeptides significantly changed in control and 10 Hz stimulated groups. pPDH was significantly enriched in the control group. (7,542 plotted phosphopeptides, cutoff fold change = 2, p = 0.05).

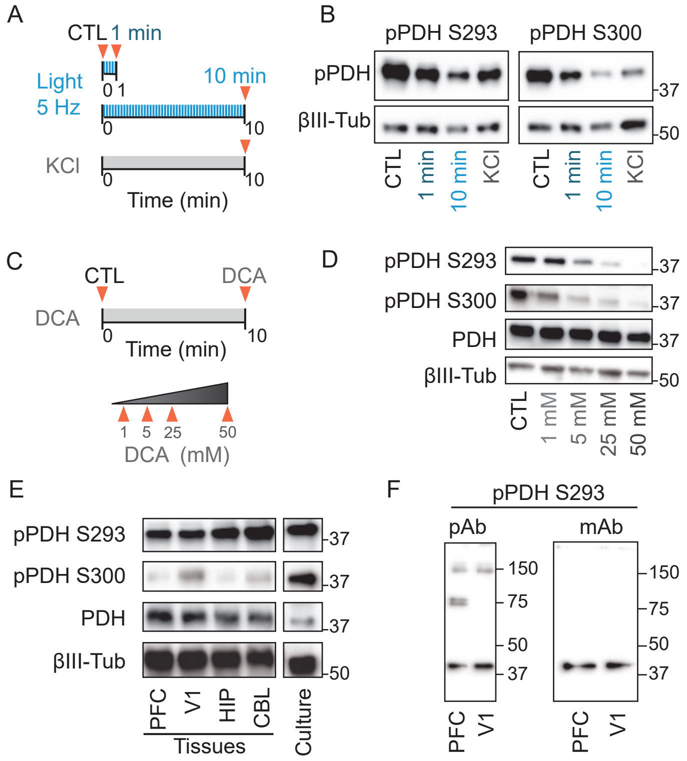


**Figure S2. Additional characterization of stimulation effects on pPDH, related to Figure 2**

(A-B) The effect of neuronal activity on pPDH. (A) Cultured cortical neurons were applied with either blue light (5 Hz) for 1 minute or 10 minutes, or KCl (50 mM) for 10 minutes. Cells were lysed at the end of stimulation (marked by red triangles) for immunoblot analysis (B).

(C-D) The effect of pyruvate dehydrogenase kinase inhibitor (dichloroacetate (DCA)) on pPDH. (C) Cultured cortical neurons were treated with DCA at different doses, and cells were lysed for immunoblot analysis (D).

(E) pPDH (S293) is the dominant phosphorylation site in vivo. Whole-cell lysates of mouse brain tissues (PFC, V1, HIP, and CBL) and cultured neurons were analyzed by immunoblotting for the phosphorylation sites of PDH.

(F) Comparison of pPDH polyclonal (Millipore AP1062) and monoclonal (Cell Signaling 37115) antibodies by immunoblot analysis of PFC and V1 whole-cell lysates.

PFC: prefrontal cortex; V1: primary visual cortex; HIP: hippocampus; CBL: cerebellum.

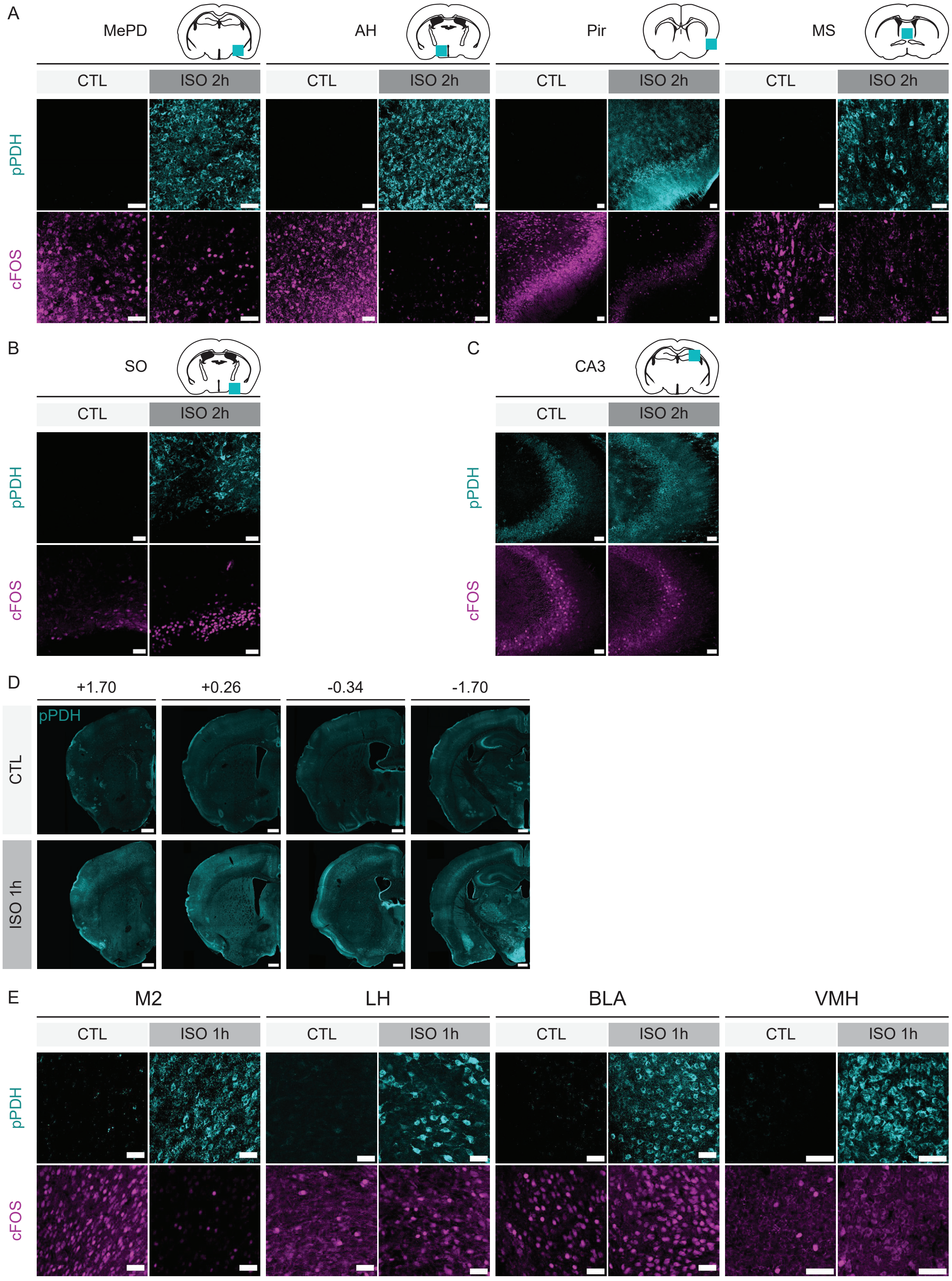

A
MePD
AH
Pir
MS
CTL
ISO 2h
pPDH
cFOS
B
SO
C
CA3
D
+1.70
+0.26
-0.34
-1.70
CTL
ISO 1h
E
M2
LH
BLA
VMH

**Figure S3. Additional characterization of pPDH in general anesthesia, related to Figure 3**

(A-C) Representative zoomed-in images of pPDH and cFOS labeling in different brain regions after 2 hours of general anesthesia.

(D-E) pPDH labeling in the whole brain after 1 hour of general anesthesia. (D) Representative images of coronal brain sections at indicated bregma positions after 1 hour of general anesthesia. (E) Representative zoomed-in images of pPDH and cFOS labeling in different brain regions. N = 3 mice for control and N = 4 mice for isoflurane treated group.

Scale bar: 50 μm in (A-C and E), 500 μm in (D).

MePD: medial amygdaloid nucleus, posterodorsal part; AH: Anterior hypothalamic area; Pir: piriform cortex; MS: medial septal nucleus; SO: supraoptic nucleus; CA3: field CA3 of hippocampus; M2: secondary motor cortex; LH: lateral hypothalamic area; BLA: basolateral amygdaloid nucleus, anterior part; VMH: ventromedial hypothalamic nucleus.

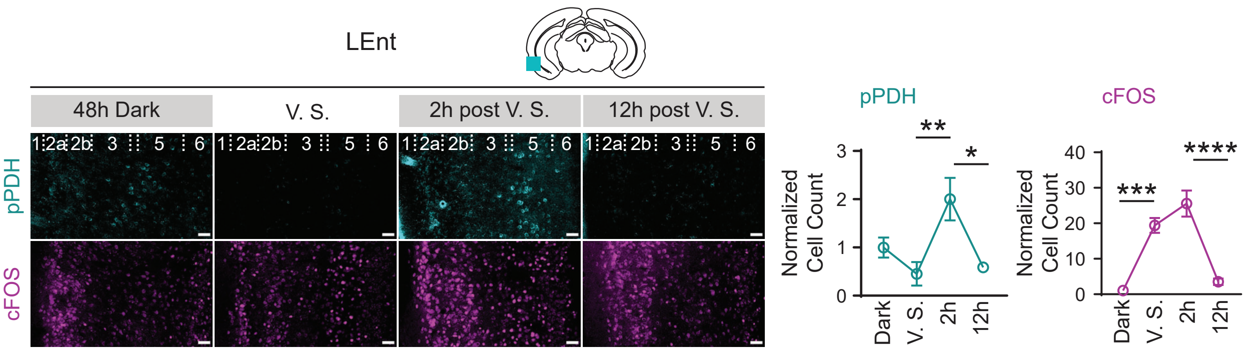


**Figure S4. Additional characterization of pPDH in visual stimuli, related to Figure 4**

Representative images of pPDH and cFOS staining in LEnt after visual stimuli. Positively stained cells in layer 5 were quantified and normalized to the 48 hour dark group. For LEnt: 48 hour dark (N = 4), V.S. (N = 4), 2 hour post V.S. (N = 4), 12 hour post V.S. (N = 4).

All values are mean ± s.e.m. Statistics determined by ordinary one-way ANOVA with Tukey's multiple comparisons test. * $p < 0.05$; ** $p < 0.01$; **** $p < 0.0001$.

Scale bar: 50 μm.

LEnt: lateral entorhinal cortex.

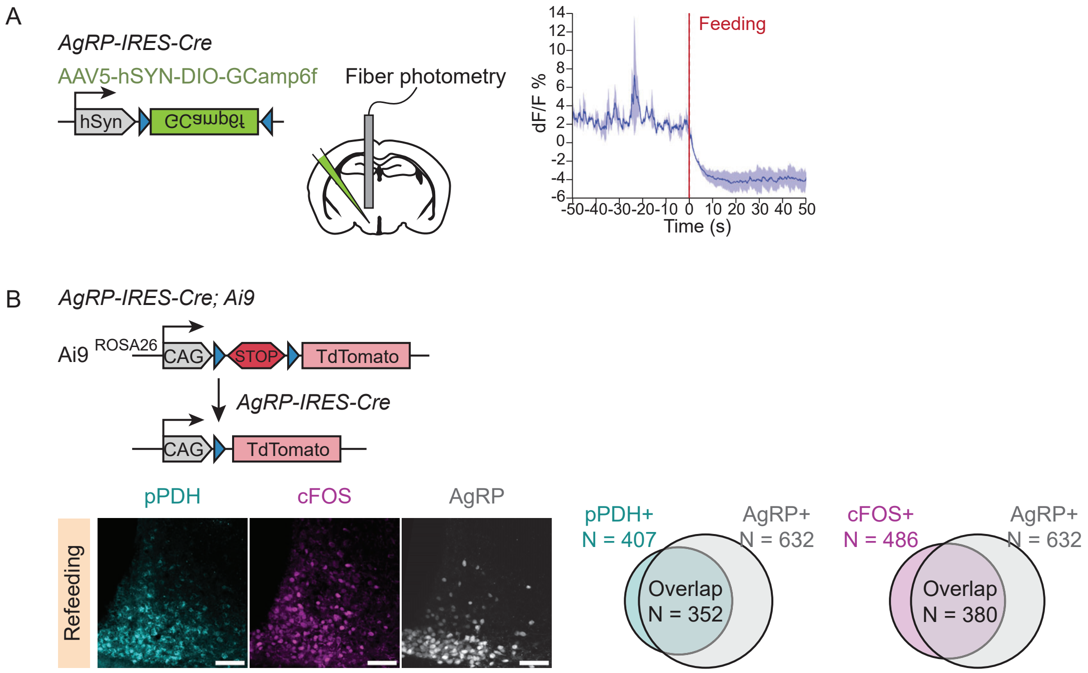


**Figure S5. Additional characterization of pPDH labeling in the ARC, related to Figure 5**

(A) Left, schematic of virus injection and fiber photometry. Right, Calcium signal from AgRP neurons in fasted mice presented with food. (N = 2 mice).

(B) Top, schematic of the genetic strategy used to generate AgRP labeling mice. Bottom, representative images of pPDH and cFOS labeling in ARC of AgRP-Ai9 mice 1 hour after feeding. pPDH and cFOS labeled cells were quantified. N = 4 mice.

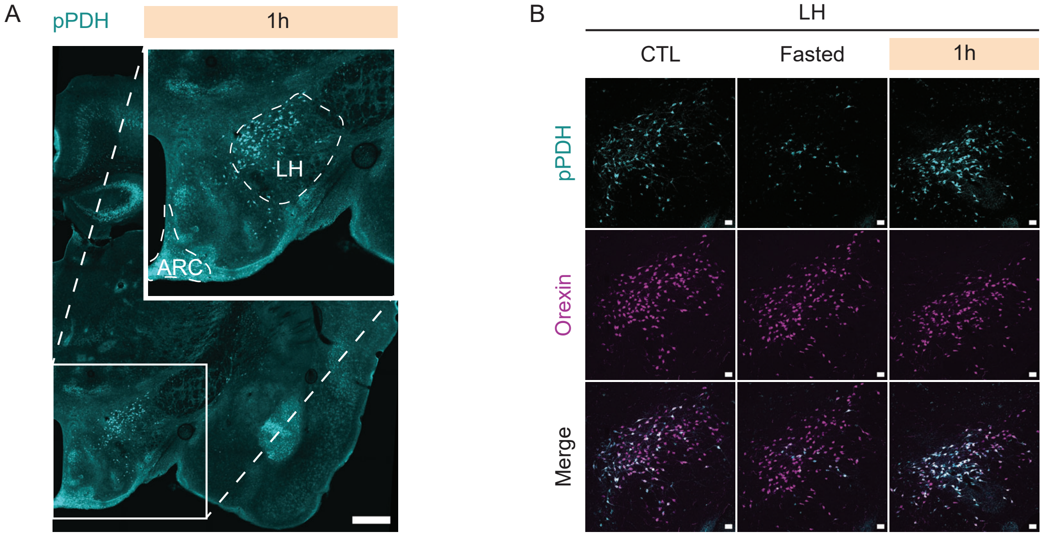


**Figure S6. Additional characterization of pPDH labeling in the LH, related to Figure 6**

(A) Representative image of brain coronal section showing pPDH labeling in ARC and LH in 1 hour refed mice.

(B) Representative images of pPDH and Orexin labeling in LH in fasting-refeeding paradigm.

Scale bar, 500 µm in (A), 50 µm in (B).

ARC: arcuate hypothalamic nucleus; LH: lateral hypothalamic area
